# Benchmarking fungal species classification using Oxford Nanopore Technologies long-read ITS metabarcodes

**DOI:** 10.1101/2025.07.02.661940

**Authors:** Abigail Graetz, Jinghang Feng, Alex Ringeri, Austin Bird, Duong Vu, Camille Truong, Benjamin Schwessinger

## Abstract

The gold standard for fungal classification has long been specimen or culture-based, however, many Fungi display simple morphological characteristics, or are unculturable by current methods. Using DNA metabarcodes, many taxa can be identified from a single sample simultaneously, but annotating these sequences with species-level taxonomy often requires more information than short-read sequences afford. Oxford Nanopore Technologies (ONT) long-read sequencing has achieved species-level resolution in metabarcoding of bacteria and invertebrates, but complex fungal taxonomy and biology has been a hurdle to the application of this technology for Fungi. In this work, we use a mock community of real ONT long-read metabarcodes from 54 fungi from the Dikarya subkingdom to extensively benchmark classification approaches to assign species-level taxonomy. We compare eight classification approaches spanning alignment and *k*-mer based algorithms, to emerging machine learning methods, and assess the sensitivity, precision, and diversity estimation of each classifier at the species level. Our results indicate that classifiers which determine informed thresholds of sequence similarity based on a provided reference database are not only more accurate at the species level, but more consistent to correct species abundance distributions, and better able to place sequences from ‘unknown’ species taxonomically closer to their true origin. We demonstrate the power of machine learning classifiers to leverage long-read metabarcodes, and their promise as emerging methods in DNA sequence classification. We present our results in the context of real-world use cases, to demonstrate that species-level taxonomic inference is achievable, precise, and reliable with ONT long-read fungal metabarcodes.

## **1.** Introduction

### 1.1 The ITS as the Fungal metabarcode

Despite their taxonomic diversity, many Fungi display simple morphological characteristics, and a large proportion are unculturable by current methods (Abarenkov et. al., 2022; Nilsson et. al., 2016; Lücking et. al., 2020). DNA-based identification using loci known as DNA barcodes (Hebert et al., 2003) has emerged as the primary method for overcoming this limitation, dramatically improving our capacity to detect and identify fungi from environmental samples (Větrovský et al., 2019; Tedersoo et al., 2022; Ji et. al., 2013; Rieker et. al., 2024; Yahr et al., 2016; Phukhamsakda et al., 2022; Abarenkov et al., 2022; Nilsson et al., 2016; Lücking et al., 2021). The formalised DNA barcode for most members of the Kingdom Fungi is the internal transcribed spacer (ITS) region of the ribosomal DNA (rDNA), chosen for its high interspecific variability and its conserved primer binding sites across Fungi (Schoch et al., 2012; Figure 1). Building on the technique of DNA barcoding, DNA metabarcoding detects and differentiates between many taxa in a single sample (Ji et al., 2013; Cristescu, 2014). The fungal ITS can also be used for this purpose, as conserved ribosomal subunit genes can be targeted with broad-spectrum primers to amplify the ITS region from many species simultaneously (Martin and Rygiewicz, 2005; Ihrmark et al., 2012; Tedersoo et al., 2008). Broadly, metabarcoding has applications in pathogen detection (Ohta et al., 2023; Huggins et al., 2024; Chen et al., 2022), ecosystem monitoring (Abrego et al., 2024; Cordier et al., 2018; Keck et al., 2018; Gorki et al., 2024), and biodiversity assessment (Taberlet et al., 2012; Rieker et al., 2024; Mikryukov et al., 2023) including microbiome characterisation (Bahram et al., 2018; The NIH Human Microbiome Project Working Group et al., 2009).

**Figure 1.**
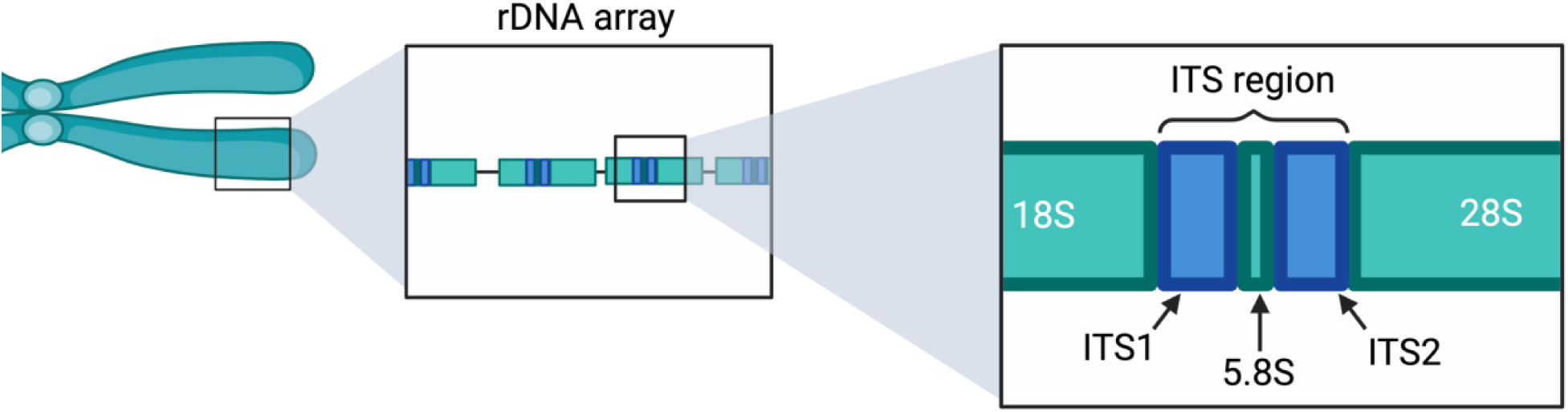
The internal transcribed spacer (ITS) is a part of the ribosomal DNA (rDNA) array. The ITS1 and ITS2 are physical spacing regions separating the genes encoding the catalytic components of the ribosome. In Eukaryotes, the rDNA is tandemly repeated, and is usually found on the arm of one chromosome. Figure created using Biorender.com.

The ITS is an imperfect barcode for Fungi, as low interspecific variation in some fungal groups prevents differentiation of closely related species (Puillandre et al., 2011; Stoekle and Thaler, 2014; Stockinger et al., 2010), yet because the ITS is a multi-copy region, there can also be high intraspecific variation (Estensmo et al., 2021; Hilário et al., 2022; Tremble et al., 2020). While some authors argue that clustering steps are important for avoiding oversplitting due to intraspecific variations (Estensmo et al., 2021; Kauserud, 2023), this can mask true sequence variation between species (Elbrecht et al., 2018; Nilsson et al., 2008; Yamamoto and Bibby, 2014) and obscure calculations of species relative abundance in community ecology studies (Larvinienko et al., 2021).

By increasing query sequence length, more comparative information can be leveraged to differentiate between species in a metabarcoding study. Short subregions of full-length metabarcodes can only confidently provide genus-level resolution (Ovaskainen et al., 2024; Matsuo et al., 2021; Irinyi et al., 2016; Johnson et al., 2019). In comparison, Oxford Nanopore Technologies (ONT) long-read sequences can span kilobases of DNA, easily containing ITS1, ITS2, and additional taxonomically relevant sequences (Lu et al., 2022; Krehenwinkel et al., 2019; Mafune et al., 2020; Ohta et al., 2023; Heeger et al., 2019; Hu et al., 2022), through to entire rDNA alleles (D’Andreano et al., 2021, Fultz et al., 2023; Wurzbacher et al., 2019), improving taxonomic resolution. ONT long-read sequencing has not been widely adopted in fungal metabarcoding largely due to its notoriously higher error rate in comparison to short-read sequencers. However, the latest ONT MinION flowcells with R10 Q20+ chemistry, along with improved basecaller models, have been demonstrated to achieve a mean per base accuracy of greater than 99%, much closer to Illumina and Pacific Biosciences sequencers (Lerminiaux et al., 2024; Sanderson et al., 2023; Srivathsan et al., 2021; Morrison et al., 2020; Cuber et al., 2023; Chang et al., 2024). With this improved sequence quality, and increased sequence length, long-read metabarcodes can improve taxonomic classification to the species level, reducing uncertainty in clinical settings, achieving finer resolution in community characterisation for more accurate understanding of microbial interactions, and refining phylogenetic placement of taxa in both fungi (Ohta et al., 2023; Yang et al., 2018; Groben et al., 2023), and other eukaryotic study systems (Huggins et al., 2024; Bludau et al., 2025; Dunthorn et al., 2014).

In a metabarcoding study, the choice of classification algorithm, and the database of reference sequences used for taxonomic annotation are two of the most critical. Many analysis approaches are optimised for use with short-read sequences, particularly for use with clustering methods. Additionally, extensive benchmarking efforts for ONT long-read metabarcoding have largely focused on bacteria, microeukaryotes, and invertebrates (Bludau et al., 2025; Huggins et al., 2024; Baloğlu et al., 2021; Curry et al., 2022; Ammer-Herrmenau et al., 2021). In this work, we aim to fill this knowledge gap by benchmarking fungal metabarcode classification at the species level using mock communities comprising real ONT long-read sequences, focusing on two main variables: classifier, and reference database.

### 1.2 Benchmarking metabarcode classification: reference databases

Reference databases can strongly influence the interpretation of metabarcoding data, due to missing taxa or mislabelled records which can lead to misclassification and misleading results. This aspect of database limitation has been thoroughly reviewed in several other publications (Hestetun et al., 2020; Dias et al., 2020; Keck et al., 2023; Kvist, 2013). Instead, our benchmarking aims to compare the application of different reference databases for specific use cases, using an in-house database as an example of a small, specific database, and the NCBI RefSeq Targeted Loci Fungi ITS database (O’Leary et al., 2016; Schoch et al., 2014) as an example of a broader reference database.

The ‘best’ reference database to use may not be the one with the most sequences, or the most species represented, particularly in the case of long-read metabarcoding. In this light, we tested two database frameworks. The first, is a highly targeted database, with reference sequences of similar length to the input query sequences. While using a larger database may be intuitive when profiling community diversity, this can reduce accuracy and confidence when species of interest are not present. Smaller, study system-specific databases are gaining popularity for diagnostic purposes (Langsiri et al., 2023; Ohta et al., 2023; Chen et al., 2022), and ecological surveys in underexplored ecosystems (Bourret et al., 2023; Philip et al., 2024) because users can be ensured that species in their system of interest will be better represented. However, for other uses, such as mycobiome profiling from environmental DNA (eDNA), publicly available reference databases such as UNITE (Abarenkov et al., 2024) or NCBI RefSeq (O’Leary et al., 2016; Schoch et al., 2014) which contain sequences for thousands of fungal species may better represent the diversity of the study system. While these databases are still incomplete, in study designs where fungal community composition is unknown, broad databases may provide better estimates of community diversity and ecology than smaller, specific databases.

### 1.3 Benchmarking metabarcode classification: classifier

Taxonomic classification of DNA sequences relies on similarity searches between query sequences provided by the user and the sequences in a reference database, to assess the likely origin of a query sequence based on pairwise sequence identity. How this similarity is assessed and the minimum similarity threshold required for confident taxonomic classification is complicated by fungal taxonomy, the length of ONT query sequences relative to database reference sequences, and classification approach. There are three general categories for per-sequence taxonomic classification (that is, without a clustering step): alignment-based approaches, *k*-mer search methods, and emerging machine-learning methods.

The most common sequence similarity approach for taxonomic classification is a search using the Basic Local Alignment Search Tool (BLAST) algorithm (Altschul et al., 1990; Camacho et al., 2009), which classifies query sequences based on their closest pairwise alignment in a reference database (Taberlet et al., 2018; Tedersoo et al., 2022). While BLAST is highly sensitive, it may not be the most suitable classification option for long ITS metabarcoding approaches (Lu et al., 2022), especially for large datasets of many sequences (Ye et al., 2019). Additionally, using fixed threshold values of local alignment similarity to delimit taxa, such as with a BLAST search, does not accurately represent true species-level boundaries of sequence similarity - a particularly important consideration in metabarcoding applications where many species may be present in a single sample (Vu et al., 2016; 2019; 2022). Dnabarcoder is a tool that has been developed to address this issue by predicting separate similarity cutoffs for different fungal groups using a reference sequence dataset, and then classifying query sequences using these predicted cutoffs with BLAST (Vu et al., 2022). Dnabarcoder’s per-species similarity cutoffs significantly improved the accuracy and specificity of sequence classification when tested on a large ITS barcode dataset (Vu et al., 2022), but have not been benchmarked for use with ONT long-reads.

Lighter alternatives to BLAST, such as minimap2 (Li, 2018; Li, 2021) and Kraken2 (Wood et al., 2019), are popular for long-read sequences. Both minimap2 and Kraken2 use minimisers (Roberts et al., 2004) as intermediate query objects to reduce computational load, but differ in classification strategy. Minimap2 uses a similar ‘seed-chain-align’ strategy to BLAST, but is designed to handle much longer alignment lengths (Li, 2018; Li, 2021). In contrast, Kraken2 generates unique sub-sequences of length *k* (*k*-mers) from the provided reference sequences at different taxonomic ranks, then uses these to classify query sequences to the lowest common ancestor (LCA) (Wood et al., 2019). Several studies have been published using these methods to classify long-read ITS amplicons (Hu et al., 2022; Lu et al., 2022; Erlandson et al., 2024; Baramidze et al., 2024), and ONT’s own analysis agent EPI2ME provides both minimap2 and Kraken2 as classifier options within the automated metabarcode analysis pipeline (Garcia et al., 2024).

Most recently, machine learning methods have been explored for sequence classification tasks. Strictly speaking, these methods do not use heuristics which rely on sequence similarity the way alignment and *k*-mer based methods do, but learn how to discriminate between classes by determining the hallmarks of Class 1 versus Class 2 during model training. For the purpose of DNA sequence classification, these discriminatory hallmarks may be sequence variations, which allow the model to assign a sequence to the ‘class’ of a species. In this sense, while the model is technically assessing the similarity between a query and reference sequence, the process is not as intuitive as for pairwise alignment methods. Deep neural networks, such as convolutional neural networks (CNN), are a popular approach for ML-based DNA sequence classification because they inherently parallelise, and are well-suited to complex data (Gunasekaran et al., 2021; Soliman et al., 2021; Das et al., 2023; Vu et al., 2020).

For long ITS sequences which amplify significant regions of the conserved 18S and 28S ribosomal subunits, the proportion of the sequence which contains interspecific variation may be relatively small. Transformer (Vaswani et al., 2017) is an attention-based model which greatly improved the classification accuracy of DNA sequences, as the model can focus on only the variable part of the whole sequence (Clauwaert and Waegeman, 2022; Choi and Lee, 2023; Gokhale et al., 2023; Khan and Lee, 2023; Sadad et al., 2023). Large, complex, transformer-based models have been developed in Natural Language Processing (NLP), such as BERT (Bidirectional Encoder Representations from Transformers, Devlin et al., 2019), which has a significant improvement in how machines understand human language. MycoAI (Romeijn et al., 2024) is an example of an adaptation of this kind of model specifically for processing gene sequences, which regard the sequences as sentences in natural language, and demonstrated improved accuracy in comparison to alignment-based methods (Ji et al., 2021; Mock et al., 2022; Gwak and Rho, 2022; Helaly et al., 2022). This is in part because of the more nuanced way that these types of models assess sequence similarity, in comparison to pairwise alignment methods.

The Emu algorithm is one of few hybrid methods which exist for classifying metabarcodes (Curry et al., 2022). Initial taxon probability distributions are estimated by aligning query sequences to database sequences with minimap2, then an expectation-maximisation (EM) algorithm (Do and Batzoglou, 2008; Ceppellini et al., 1955) applied to reduce noise originating from the ONT error profile and prevent overestimation of species diversity. While Emu was intended for, and tested on, bacterial 16S metabarcodes, the algorithm can be applied to ITS metabarcodes with the provision of an appropriately formatted database (Nagpal et al., 2024; Erlandson et al., 2024).

Given the recent emergence of ML methods, there is no comprehensive benchmark applying these methods to long-read fungal metabarcodes, or comparing alignment and *k*-mer based techniques. To provide a thorough assessment of classification methodologies, we use real ONT long-read sequences from 54 diverse fungal taxa to benchmark taxonomic assignment strategies for long ITS sequences. We present our results in the context of two broad use cases: biodiversity assessment, where accurately profiling diversity and relative abundance is a priority; and a diagnostic scenario, where understanding detection limits, false positive rate and false negative rate are a priority. We demonstrate high precision of identification to the species level, and accurate reproduction of species abundance distributions with ONT long-read ITS metabarcode sequences. Classification methods can be robust to missing species in reference databases, particularly those approaches which determine informed per-species thresholds of sequence similarity from the reference database prior to classification. Our results also illustrate the power of machine-learning methods to leverage the full length of the ITS metabarcode to discriminate between fungal species, with these models outperforming alignment and *k*-mer based methods when provided with sufficient training data.

## 2. Methods and Materials

### 2.1 Benchmarking dataset

We sequenced long-read ITS amplicons from 54 fungal taxa (Supplementary File S1), representing 28 distinct Families from within the Kingdom Fungi.

Initially, 60 fungal species underwent DNA extraction independently from pure culture material (spore or mycelial), and the full ITS region amplified by PCR using primers NS5 (forward) and LR6 (reverse) (White et. al., 1990). Amplicons were prepared for sequencing using the ONT Native Barcoding Kit (SQK-NBD114.96) and sequenced on an R10.4 MinION flowcell via the MinKNOW graphical user interface. Full protocols for PCR and library preparation are available in Supplementary File S2.

All sequences were basecalled and demultiplexed with Guppy (version 6.4.2, super-high accuracy model v3.5.1) and adapters removed with the --trim_adapters option. Primer sequences were removed using porechop (v0.2.4, Wick et. al., 2017), and sequences filtered to minimum length of 300 bp, maximum length of 6000 bp, and minimum average Phred score of 15 (i.e., ∼95% accuracy) or 17 (i.e., ∼98% accuracy) using chopper (v0.7.0, De Coster and Rademakers, 2023) (Supplementary File S3). After sequence quality control, six species were removed from the mock community due to low quality, and/or indistinguishable ITS sequences (100% nucleotide identity with another species). Quality controlled sequences were subsampled to create *in silico* mock communities of known species abundances, as detailed in Supplementary File S1. Note that abundance distributions were created for 60 species, even though only 54 of these were used.

### 2.2 Mock Community 1: Fungal classification in evenly abundant communities

We first used the quality-controlled sequences to create an evenly distributed *in silico* mock community. To generate *in silico* Mock Community 1 (Mock 1), 2,000 sequences from each single-species FASTQ file were randomly subsampled without replacement into a single FASTQ file, to create a community of 54 taxa with even abundance.

### 2.3 Mock Community 2: Fungal classification in unevenly abundant communities

In a community composition survey, taxa are not evenly distributed. Microbiome data, for example, are known to require various transformation techniques to address issues of data sparsity (i.e., zero-inflation), dispersion, and compositionality (Thorsen et. al., 2016; McKnight et. al., 2019; Karwowska et. al., 2025).

To generate *in silico* Mock Community 2 (Mock 2), sequence abundance was normally distributed around a mean of 2,000 sequences/sample in R, which was used to determine the number of sequences to be subsampled for each species. Sequences were randomly subsampled without replacement from each quality controlled single-species FASTQ files to the desired relative abundance, as detailed in Supplementary File S1.

### 2.4 Mock Community 3

Detection of a low abundance, known pathogen from close relatives. While some pathogens may establish in high abundance and with specific phenotypes, others may cause disease at low abundance in the host, or may not be easily isolated, limiting their relative abundance in a DNA sample. To assess the ability of each classification approach to detect a low abundance pathogenic fungus from a mixed sample of close relatives, *in silico* Mock Community 3 (Mock 3) was generated, with *Candida tropicalis* as the target pathogen. Relative abundance of *C. tropicalis* for this experiment was 7.2% of *Candida* group sequences (0.77% of total Mock 3 sequences). Abundance of non-*Candida* taxa was distributed around a mean of 2,000 sequences/taxon (+/-1,200), simulated in R. Sequences were randomly subsampled without replacement from each quality controlled single-species FASTQ file to the desired relative abundance, as detailed in Supplementary File S1.

### 2.5 Classification methodologies

For all experiments, the following classification methodologies were tested for the three mock communities: aodp (Zahariev et. al., 2018), Dnabarcoder (Vu et. al., 2022), Emu (Curry et. al., 2022), Kraken2 (Wood et. al., 2019), minimap2 (Li, 2018; Li, 2021), the CNN model published by Vu et. al., (2020), referred to as the ‘vtd’ classifier, and the CNN and BERT models of mycoAI (Romeijn et. al., 2024). These methodologies represent alignment-based, *k*-mer based, and machine learning methods, however we broadly divided these into alignment-based (including *k*-mer methods), and machine learning (ML)-based approaches. Detailed descriptions of run conditions, including code, are available at: https://github.com/AbiGraetz/Nanopore-ITS-Benchmarking.

### 2.6 Databases for alignment-based classifiers

For alignment-based methods, each mock community was classified against two databases. The first is referred to as the ‘Gold Standard Database’ (GSD), which consisted of consensus sequences generated from the 54 taxa used to create mock communities. Supplementary File S4 details the consensus building approach. All taxa present in the mock communities were represented in the GSD, with a high-quality consensus sequence expected to be highly similar in both length, and nucleotide sequence, to the query sequences for that taxon.

The second database used for alignment-based methods was the NCBI RefSeq Targeted Loci Fungi ITS database (O’Leary et al., 2016; Schoch et al., 2014, release 226, downloaded November 2024). This database contains sequence records for over 19,600 fungal species, largely from Type material (Goldfarb et al., 2025). NCBI RefSeq ITS records only 35 of the 54 taxa in our mock communities, and contains on average much shorter sequences than the query amplicons. Note that we refer to this as the ‘NCBI database’, or ‘NCBI RefSeq’ for brevity, unless otherwise specified.

### 2.7 Training sets for machine-learning classifiers

For ML-based methods, training sets were required to train models prior to classification. For the training set emulating the GSD, we sampled 2,200 sequences per GSD species without replacement, and without crossover between the training and testing datasets, from quality controlled per-species FASTQ files. Because this training set is comprised of the same sequences which created the GSD consensus sequences, it is also referred to as the GSD training set.

To provide a suitable training set for comparison to the NCBI RefSeq database, we took the reference sequences for the 35 species from our mock communities which appear in this version of NCBI RefSeq database (release 226, downloaded November 2024), and created a simulated training set, which we refer to as simNCBI. To re-construct longer ITS sequences, we searched for the 35 species appearing in the NCBI RefSeq ITS database in two other NCBI RefSeq Targeted Loci Projects: Fungi 18S (SSU) and Fungi 28S (LSU) (Supplementary Table T1). 18S and 28S sequences were downloaded where available, and long ITS regions were reconstructed in Geneious Prime (https://www.geneious.com, v. 2024-10-30), by mapping NCBI RefSeq sequences to the GSD consensus sequences using the Geneious aligner with default settings. Long ITS reference sequences for the 35 species were imported into RStudio (R Core Team, 2021; Posit team, 2025), and copied 2,200 times, the same number of sequences per species as for the GSD training set.

To introduce variation to the simNCBI training set, we simulated mutations *in silico* to represent ‘biological variation’ and ‘sequencing error’ in R. 1% of sequences per species were randomly selected, and six positions defined to introduce small indel-like mutations of controlled size. These indel-like mutations were consistent within a species, but different across species.

As the sequences for the GSD training set were filtered to minimum Q score of 15, which represents 96.8% sequence accuracy, we distributed sequencing error values over a normal distribution bounded between 0% and 3.16%, +/-2 standard deviations from the mean. For each sequence per species, percentage error was converted to the number of positions to mutate based on sequence length. Substitution mutations were introduced at randomly selected positions per sequence, with an equally weighted probability of selecting one of the four IUPAC nucleotide letter codes for each substitution event. This process produced a dataset of sequences which approximately represent simplified, but realistic, distributions of sequence variation.

### 2.8 Classifier performance assessment

After classification, the number of true positive, false positive and false negative calls were determined at the species level for each species in the mock community. The precision (Equation 1), recall rate (Equation 2), and F1 score (Equation 3) were calculated for all combinations of mock community, classifier, and database, to assess performance of each classifier:

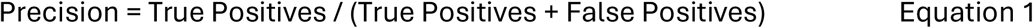

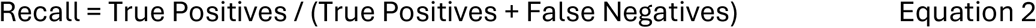

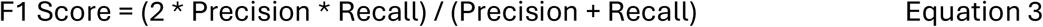

The relative contribution of experimental factors ‘Classifier’, ‘Database’, ‘Species’ and ‘Minimum Sequence Quality’ to F1 score variation was assessed with random forest regression simulations in R, using the package randomForest (Liaw and Wiener, 2002).

Because the ground truth composition of mock communities was known, Bhattacharyya distance (Aherne et. al., 1998) was used to quantify the difference between ground truth, and experimentally determined taxon abundance distributions for each mock community and database combination. Bhattacharyya distance (Equation 4) measures the degree of overlap between two probability distributions (denoted *p* and *q*).

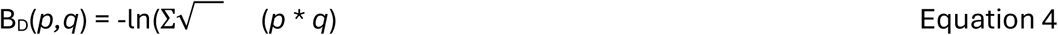

When one distribution is the ‘known’ or ‘true’ distribution, Bhattacharyya distance indicates the amount by which the observed distribution (*q*) deviates from the true distribution (*p*).

Shannon’s alpha diversity index (Shannon, 1948) was calculated for all mock communities using species-level abundances. To better compare between classifiers, the delta between the Shannon index for the ground truth community composition and the observed community composition was determined.

Mock 1 and Mock 2 datasets were also split into five equal subsamples to produce technical replicates, and classified against the GSD and NCBI/simNCBI databases using the same parameters as for the full datasets. Replicate results were used to calculate the mean, standard deviation (SD) and 95% confidence intervals (CIs) for both Bhattacharyya distance and delta Shannon index. Per-classifier means were compared within levels of Database by pairwise Dunn’s tests with Šidák correction for multiple comparisons using the function dunnTest from the FSA package in RStudio (Ogle et al., 2025).

### 2.9 Assessing classifier behaviour with unseen species

The NCBI database used for alignment-based methods only contained species-level records for 35 of the species in our mock communities, after resolving synonymous nomenclature. This left 19 species for which there was not a correct species-level classification. To assess how classifiers handled incomplete reference databases, for each of the 19 ‘unknown’ species, we found the ‘top hit’, defined as the database taxon the most input sequences had been classified to for a given species.

We scored each classifier based on the number of taxonomic ranks it took to find a common taxonomic assignment between the correct species, and the taxonomy of the top hit. We refer to this distance as ‘closeness score’, where a closeness score of one indicates genus-level match and six indicates Kingdom-level match.

To design a similar experiment for machine-learning methods, we created a new training set from the GSD sequences which only contained the species from our mock communities which appeared in the NCBI database, using 2,200 sequences per species. We trained the three ML classifiers on this truncated training set, and used the remaining 19 species which do not appear in NCBI RefSeq as the test set. We then calculated the closeness score for the top hit for each species, for each classifier.

To facilitate comparison between these two datasets (ML and non-ML classifiers), we found the lowest possible closeness score for each unknown taxon – that is, the lowest common taxonomic rank between the mock species, and its closest relative in the training set or database used to assign taxonomy. We could then calculate a delta value between the ‘predicted’ and ‘observed’ closeness scores. Differences between mean delta closeness scores by classifier were statistically tested by Dunn’s tests with Šidák correction for multiple comparisons.

Cumulative link mixed models (CLMMs) were used to model the variation in closeness score for alignment-based classifiers, using the R package ordinal (Christensen, 2023; v.2024.12-4.1). The number of sequences classified per top hit was scaled. For comparison, ordinal logistic regression models (OLRs) were defined using the package MASS (Venables and Ripley, 2002) in R to assess the significance of incorporating the factor Species as a random effect, as OLRs do not allow for random effects. NLME models were also tested, using the nlme function from the package nlme (Pinheiro et al., 2025; v. 3.1-168) in R. Model fitness was assessed using the Akaike information criterion (AIC) (Akaike, 1973).

### 2.10 Determining classifier minimum detection limits

We determined the minimum number of sequences required to detect all 54 species in the mock community, given the output of classifier *X*, which has a certain deviation from the ground truth species abundance distribution. We used Monte Carlo simulations (Metropolis and Ulam, 1949) to estimate the probability of meeting abundance threshold conditions when subsampling *n* sequences between 1 and the total number of sequences in the mock community, accounting for unclassified sequences. We then used a binary search to determine the minimum sequencing depth at which the estimated probability met a threshold of 95% probability that all species will be detected at a given abundance – either >0 sequences per species, or minimum 109 sequences per species, which represents 0.1% of total input sequences. The function was run 100 times, each with a different random seed, and the mean number of sequences required to meet minimum thresholds calculated for each classifier. Classification results using the GSD were used for Mock 1 and Mock 2 communities to generate minimum sequencing depths at each threshold. Ground truth species abundances were also used to simulate minimum sequencing depth required for an idealised classifier, as a comparison.

## 3. Results

### 3.1 *In silico* mock communities for benchmarking fungal ITS classification

We assessed the ability of several classification methodologies to accurately identify fungal taxa and characterise sample communities from ONT long-read ITS metabarcode sequences. Three mock communities were created *in silico* from individually sequenced fungal ITS ONT long-read sequences, to address different use cases during benchmarking (Table 1): an evenly distributed community (Mock 1), an unevenly distributed community (Mock 2) and a community with a low-abundance target pathogen (Mock 3). Each taxon was represented by independently generated DNA sequences to remove biases introduced by unequal extraction, or uneven amplification from a mixed sample. Because the focus of this work was to determine the influence of various classification strategies on species delimitation in mixed samples, the influences of these biases is not within the scope of this study (see Tedersoo and Lindahl, 2016; Bellemain et al., 2010; Feinstein et al., 2009; and Frau et al., 2019, for studies on these topics). We focused on reporting species-level analyses, because improved taxonomic resolution is a proposed technical improvement of using ONT long-read sequences over traditional short-read methodologies.

**Table 1.** Structure and purpose of *in silico* fungal mock communities. All mock communities were constructed from long-read ITS sequences individually amplified from pure specimens to avoid the influence of extraction and/or amplification biases.

|  |  |  |  |
| --- | --- | --- | --- |
|  | Mock 1 | Mock 2 | Mock 3 |
| Number of species | 54 species. | 54 species. | 54 species; diagnostic analysis focused on only 9 species. |
| Distribution of species | Even, 2000 sequences/species. | Uneven, normally distributed between 1,090 sequences, and 4,179 sequences per species. | Uneven, normally distributed with low-abundance target pathogen (0.77% of total mock community sequences, 7.2% of focus group sequences). |
| Purpose | Idealised metabarcoding study, no influence of rare or dominant species. | Simplified metabarcoding study, with low and high abundance species. | Emulating a diagnostic setting with a very low abundance pathogen. |

While we refer to some taxa in the mock community as *Candida* species for brevity, and all are members of the *Candida* group, these species represent members from several clades within *Candida* (Kidd et al., 2023). We manually verified updated, or synonymous nomenclature for all mock community taxa by careful sequence analysis and comparison to the latest literature.

Each mock community was classified against reference sequences from two databases: an in-house Gold Standard Database (GSD), and the NCBI RefSeq Targeted Loci Fungi ITS database (O’Leary et. al., 2016; Schoch et. al., 2014, release 226, downloaded November 2024) (Table 2). We used the GSD as an example of a small, but specific database, which are seeing increasing use in diagnostic cases, and biodiversity surveys in ecosystems where species richness is low. It also represents a system in which we can understand classifier performance without the confounding effects of reference database quality, such as missing records. In contrast, we used the NCBI RefSeq database as an example of a more comprehensive, publicly available reference database which may be used in biodiversity surveys in ecosystems where species richness is high.

**Table 2.** Databases used to assign taxonomic identities to long-read ITS sequences in our benchmarking work.

|  |  |  |  |  |
| --- | --- | --- | --- | --- |
|  | Gold Standard database (GSD) | NCBI RefSeq Targeted Loci ITS database (NCBI) | Gold Standard training set (GSD) | Simulated NCBI RefSeq Targeted Loci ITS training set (simNCBI) |
| Total sequence records | 54 | 78,814 | 118,800 | 77,000 |
| Total species represented | 54 | 17,536 | 54 | 35 |
| Mock community species represented | 54 | 35 | 54 | 35 |
| Mean sequence length | 2,329 bp | 592.4 bp | 2135.15 bp | 2714.29 bp |

### 3.2 Classification methodologies perform comparably well under idealised conditions

We assessed the performance of eight classifiers: aodp (Zahariev et al., 2018), dnabarcoder (Vu et al., 2022), Emu (Curry et al., 2022), Kraken2 (Wood et al., 2019), minimap2 (Li, 2018; Li, 2021), the CNN model published by Vu et al. (2020), referred to as the ‘vtd’ classifier, and the CNN and BERT models from mycoAI (Romeijn et al., 2024) (Table 3). We first tested these classifiers under idealised conditions: using a 54 species mock community where all species are evenly abundant, referred to as Mock 1 (Figure 2A), and classifying against a database (or training set, for ML methods) where each mock community species is represented by reference sequences of similar length to the query sequences.

**Table 3.** Eight classification approaches benchmarked for species-level taxonomic classification of long-read fungal ITS metabarcodes.

|  |  |  |  |  |
| --- | --- | --- | --- | --- |
| Method | Class | Reference type | ML strategy | Citation |
| minimap2 | Alignment | Database | None | Li, 2018; Li, 2021 |
| aodp | *k*-mer | Database | None | Zahariev, Chen, Visagie and Lévesque, 2018 |
| Kraken2 | *k*-mer | Database | None | Wood, Lu and Langmead, 2019 |
| Dnabarcoder | Alignment | Database | None | Vu, Nilsson and Verkley, 2022 |
| Emu | Alignment + machine learning | Database | Expectation–maximization | Curry et al., 2022 |
| vtd | Machine learning | Training set | Convolutional neural network | Vu, Goenewald and Verkley, 2020 |
| MycoAI CNN | Machine learning | Training set | Convolutional neural network | Romeijn, Bernatavicius and Vu, 2024 |
| MycoAI BERT | Machine learning | Training set | Transformer | Romeijn, Bernatavicius and Vu, 2024 |

**Figure 2.**
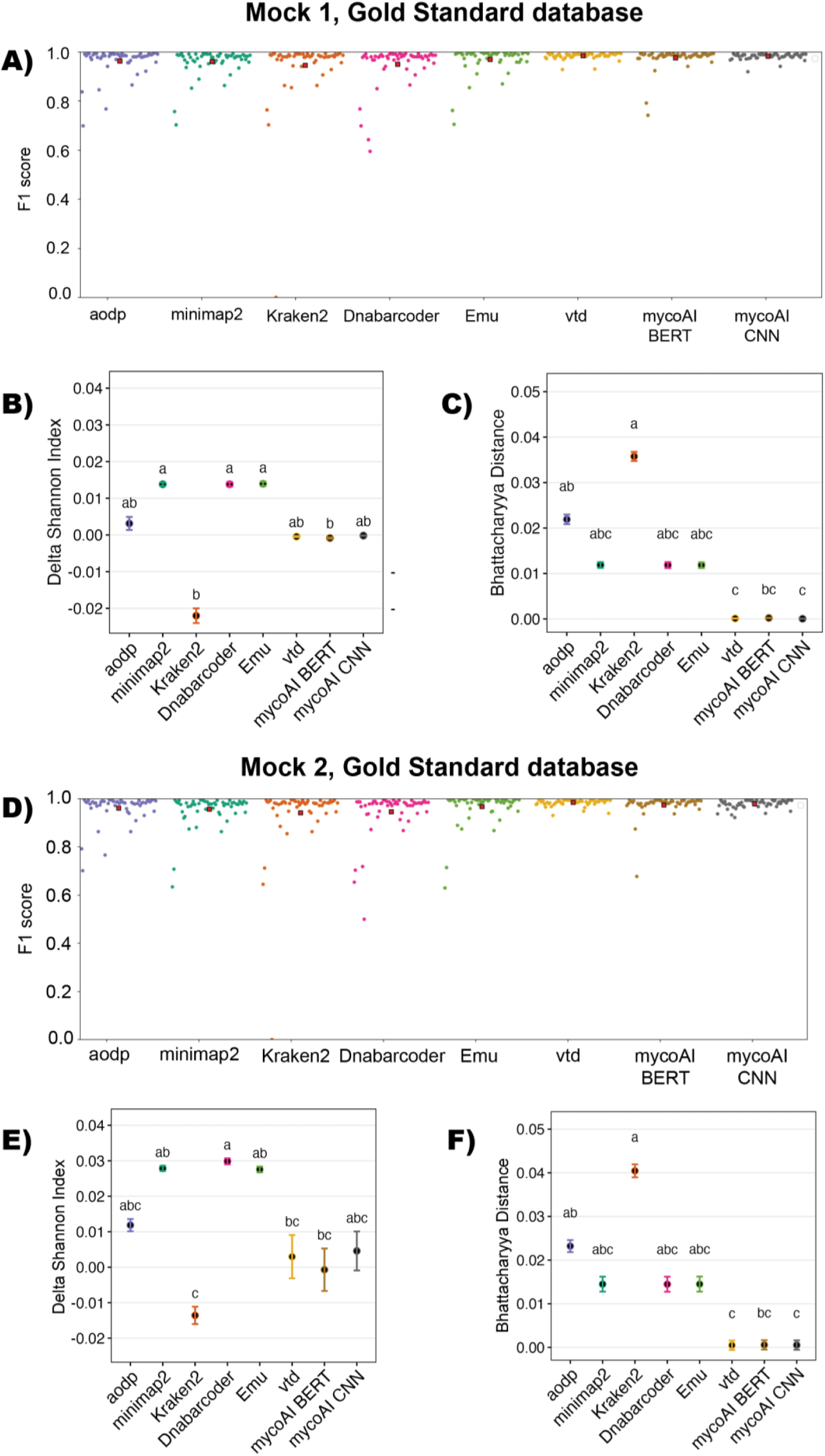
All classification approaches demonstrate high performance when tested with Mock 1 (A-C) and Mock 2 (D-F) combined with the Gold Standard Database. F1 score for Mock 1 (A) and Mock 2 (D) indicates low false positives and false negatives using this database for all classifiers tested. Machine learning methods vtd, mycoAI BERT and mycoAI CNN best reproduced community composition measured by Bhattacharyya distance and delta Shannon index in evenly distributed (B, C) and unevenly distributed (E, F) mock communities. CLD letters indicate significant pairwise differences measured by Dunn’s tests with Šidák correction for multiple comparisons (p < 0.05).

For evenly distributed Mock1 communities, mean F1 score was consistently above 0.9 for all classifiers when using the GSD as a reference, with the exception of Kraken2 (0.857) (Figure 2A), indicating that most classifiers limited both false positives and false negatives under idealised conditions. The two CNN models, ‘vtd’ and mycoAI CNN, had the largest F1 score (0.985 and 0.983 respectively), followed by mycoAI BERT (0.979) and Emu (0.975). *Aspergillus flavus* was consistently mis-identified by alignment-based approaches at the species level, despite being correctly identified at the genus level by all approaches. This is consistent with previous studies which have shown *A. flavus* to be a particularly difficult species to delineate using ITS barcoding, often only resolved to the section level (that is, above species, but below genus) (Lee and Yamamoto, 2015; Steenwyk et al., 2024).

We chose Shannon index (Shannon, 1948) to assess community alpha diversity profiles produced by each classifier, to avoid introducing bias due to rare species in unevenly distributed communities, as Simpson’s index focuses on the dominance of species (Simpson, 1949). We calculated the delta value between the observed Shannon index for a given classifier, and the Shannon index calculated from the known ground truth abundance of the input mock community, to make more balanced comparisons between classifiers. For Mock1 communities classified against the GSD, alignment-based classifiers overestimated community alpha diversity in comparison to the expected ground truth value (Figure 2B), except for Kraken2, which underestimated alpha diversity by a very small margin (-0.022). The classifiers which best reproduced Mock 1 community composition were the three ML methods, vtd, mycoAI CNN and mycoAI BERT when trained on the GSD training set.

We also used Bhattacharyya distance to capture the distance between the ground truth community abundance distribution, and the observed distributions produced by each classifier. For Mock 1, ML methods had the smallest Bhattacharyya distance (Figure 2C), indicating that the output community composition was most similar to the ground truth species distribution. In particular, the two CNN methods, ‘vtd’ and mycoAI CNN, were statistically distinct from *k-*mer methods Kraken2 and aodp (Dunn’s test with Šidák correction for multiple comparisons, p < 0.05; Figure 2C).

### 3.3 Unevenly distributed species abundances influence classifier performance

We created Mock 2, a mock community of the same 54 species unevenly distributed, to emulate a more realistic eDNA metabarcoding survey.

The machine learning methods ‘vtd’, mycoAI CNN and mycoAI BERT continued to best limit false positives and false negatives, producing the highest F1 scores when classifying the unevenly distributed Mock 2 community against the GSD (0.988, 0.986 and 0.988 respectively) (Figure 2D). ML classifiers also outperformed alignment- and *k*-mer-based classifiers in terms of delta Shannon index (Figure 2E), though were more variable across technical replicates. Bhattacharyya distance to the true community abundance distribution was also lowest for the three ML methods (Figure 2F).

In comparison to Mock 1, delta Shannon index results for Mock 2 were more positive in value for all classifiers except for mycoAI BERT. Because the GSD contains only the 54 species which were present in the mock community, and therefore the community richness could not be overestimated, this indicates that classifiers tended to overestimate the abundance of rare species, leading to increased community evenness, which increases the value of the Shannon index.

### 3.4 Dnabarcoder, Kraken2 outperform other classifiers when using a broad reference database

We used the NCBI RefSeq Targeted Loci Fungi ITS database as an example of researchers’ usage in a real metabarcoding experiment. We limited our calculations of precision, recall and F1 score to only those species where a true positive classification existed at the species level, which was 35 of our 54 mock community species. No classifier achieved a mean F1 score above 0.8 (Figure 3A) when classifying Mock 1 or Mock 2 communities. The aodp classifier did not classify any mock community sequences to the species level using the NCBI database, and so was excluded from further analyses. The other *k*-mer method tested, Kraken2, had the highest mean F1 score of tested classifiers when using the NCBI database, followed closely by Dnabarcoder.

**Figure 3.**
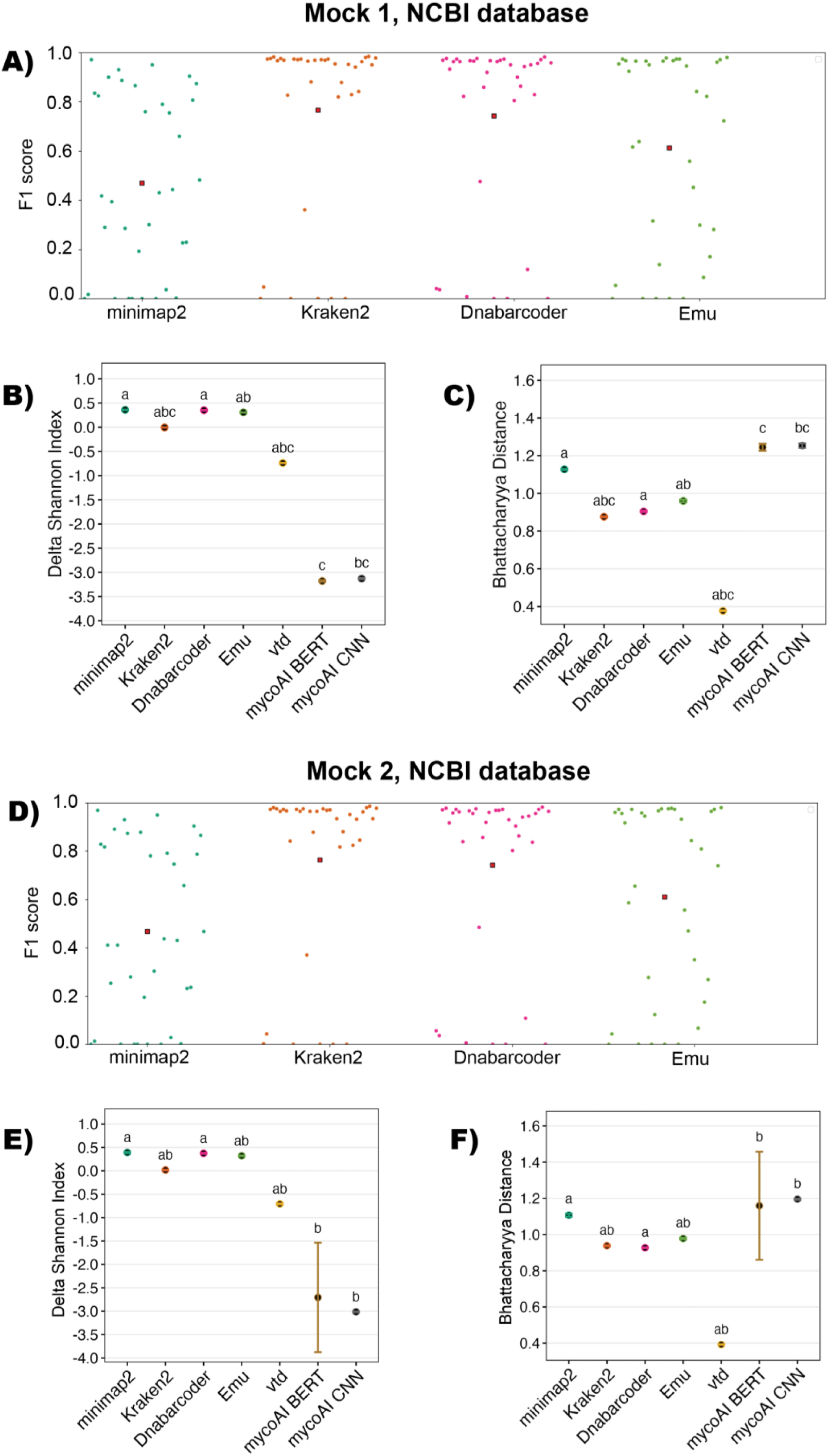
Kraken2 and Dnabarcoder outperform other classifiers when tested with Mock 1 (A-C) and Mock 2 (D-F), and the NCBI RefSeq ITS database. F1 score was highly variable for alignment-based classifiers for both mock communities (A, D). Machine learning methods did not achieve an F1 score greater than 0.5 (Supplementary Figure 2), attributable to many false negatives, as observed by low Shannon alpha diversity index values indicating underestimation of species diversity. For both even and unevenly distributed communities, Kraken2, Dnabarcoder and Emu consistently performed well based on all metrics tested. CLD letters indicate significant pairwise comparisons detected by Dunn’s tests with Šidák correction for multiple comparisons (p < 0.05).

We simulated a training set for the machine learning classifiers using the NCBI sequences as a starting point, because the NCBI RefSeq database provides only a single reference sequence per species, which is insufficient training data for ML classifiers. We refer to this as the ‘simNCBI’ database. For both Mock1 and Mock2 communities, ML methods performed poorly in comparison to their previously high precision and recall rates when trained on the GSD, achieving a mean F1 score of <0.4 in all cases (Supplementary File S5). In line with our observations for GSD classified data, this lower F1 score was largely influenced by high false negatives, which is further corroborated by ML methods underestimating community alpha diversity (Figure 3B, 3E).

We again used Bhattacharyya distance to determine the distance between ground truth and observed species abundance distributions, and delta Shannon index to determine the degree to which classifiers over- or underestimated community alpha diversity. The ‘vtd’ CNN classifier, outperformed other classification methodologies (Figure 3C, 3F) with an empirically lower Bhattacharyya distance value for both Mock 1 and Mock 2, however this value was not significantly distinct from other classifiers after p value correction for multiple comparisons. Despite having the lowest Bhattacharyya distance, the ‘vtd’ classifier underestimated community alpha diversity (Figure 3B, 3E), as did both other ML classification methods. This suggests that the ‘vtd’ classifier had a consistent misclassification error for all species, producing a similarly shaped abundance distribution, hence the low Bhattacharyya distance, but underestimating the true species abundances, resulting in a negative delta Shannon index and low mean F1 score. While alignment-based methods had larger Bhattacharyya distances and generally overestimated alpha diversity, Kraken2 had the smallest delta Shannon index value for all classifiers tested.

### 3.5 Species and Classifier jointly determine F1 Score variance when Database is controlled

As F1 score is a compound metric calculated from both recall and precision, it considers both the false positive, and false negative calls for a given class. Classifier recall was much more variable than precision for both evenly and unevenly distributed communities, when comparing by classifier using the NCBI database (Supplementary File S6). This indicates that our assessment of classifier performance was more sensitive to false negatives (accounted for in the recall calculation) than false positives (accounted for in the precision calculation).

We modelled the impact of experimental parameters on the F1 score, to determine the greatest source of false negatives. In our mock communities, there are five experimental factors which could be driving variation in F1 score – Classifier, Database, Mock Community, Species, and Minimum Sequence Quality. We used random forest models (Breiman, 2001) to compute the relative importance of the experimental factors to variance in F1 score calculated for a given input species (Figure 4).

**Figure 4.**
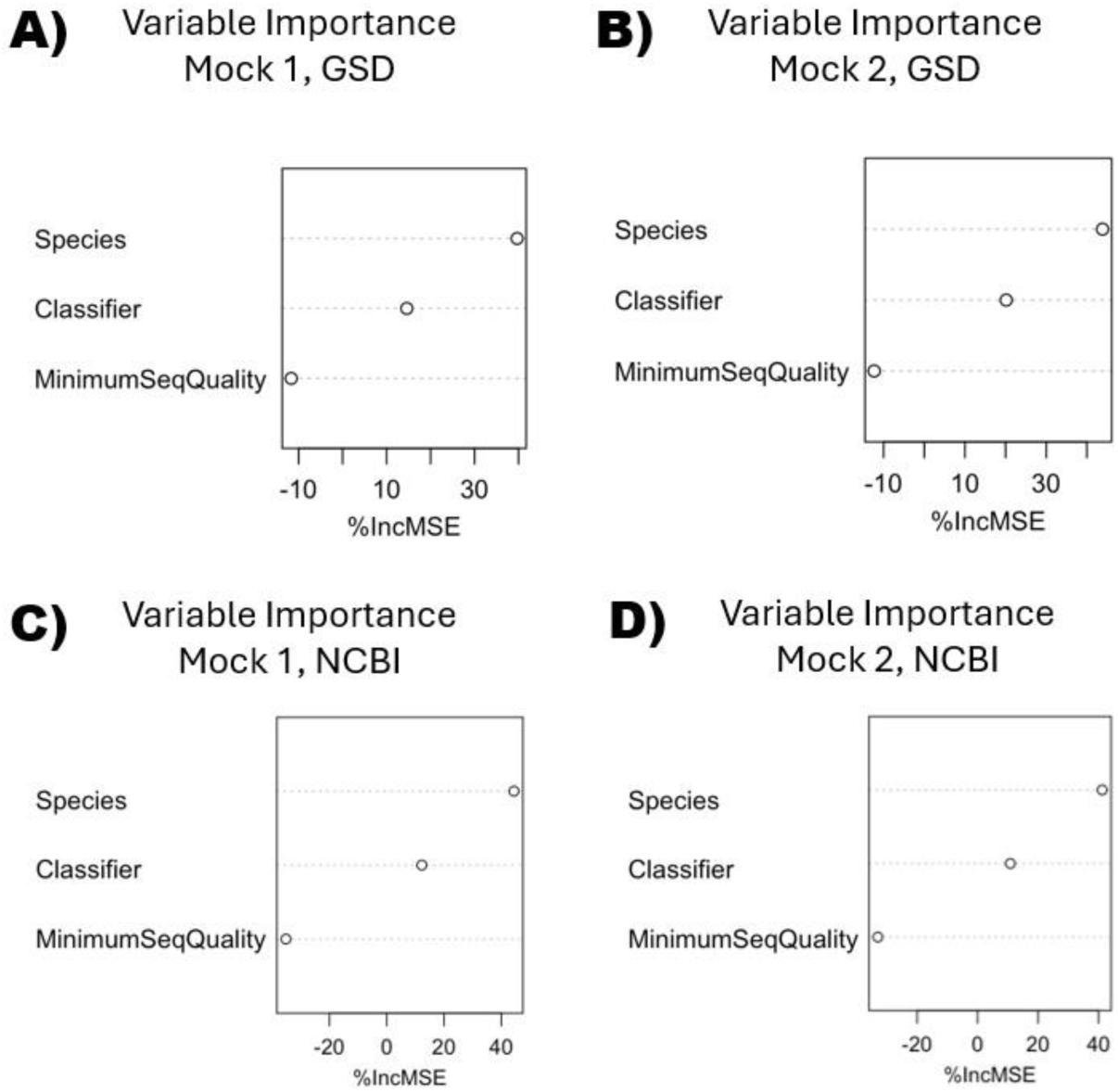
Input mock community species is a more important variable in determining F1 score than classification approach or minimum sequence quality. Random forest regression models of F1 score variance indicate that percentage increased mean squared error (%IncMSE) is highest for the factor “Species” for the evenly distributed Mock 1 community classified against the GSD **(A)**, the unevenly distributed Mock 2 community classified against the GSD **(B)**, Mock 1 classified against the NCBI RefSeq database **(C)** and Mock 2 classified against the NCBI RefSeq database **(D)**. While the factor “Classifier” was also an important variable in all models, “MinimumSeqQuality” was not important, indicated by its negative %IncMSE value.

When all factors were considered within a single mock community (Mock1 or Mock2 separately), the model captured >70% of total variance, and “MinSeqQual” was the only factor which had no importance to variance in F1 score (%IncMSE = -32.6 for Mock1, and – 34.5 for Mock2). When considering results for GSD-classified data alone, “Species” was a more important factor than “Classifier” in determining F1 score variance for both Mock1 and Mock2 (Figure 4A, 4B). This reinforces that while there is a significant effect of classifier on F1 score, different classification approaches perform comparably well when presented with a high-quality reference database, and more variation is dependent on the particular input species being considered.

Because the ML methods trained on the simNCBI database had significantly lower F1 scores per species than alignment-based methods classified against NCBI, the random forest models simulated for NCBI data excluded these results to eliminate the effect of outliers. When the F1 score results for ‘vtd,’ mycoAI CNN and mycoAI BERT were excluded, “Species” was again more important than “Classifier” in determining F1 score value, and “MinSeqQual” continued to not influence F1 score for both Mock 1 and Mock 2 communities (Figure 4C, 4D).

### 3.6 Limiting both false positive, and false negative detections in a diagnostic simulation

We used a third *in silico* mock community (Mock 3) to assess the ability of alignment-based classifiers to identify a pathogenic target species from within a mixed community of close relatives. We emphasised the performance of each classifier to accurately determine both the taxonomy, and abundance of the target pathogen *Candida tropicalis*. We assigned taxonomy to the full 54 species dataset, but focused our analysis on the nine fungi belonging to the *Candida* group, with *C. tropicalis* sequences representing 7.2% of *Candida* group sequences, or 0.77% of total input sequences.

Classifier mean F1 scores were statistically indistinguishable when using the GSD (p > 0.05, Dunn’s test with Šidák correction for multiple comparisons), and all mean F1 score values were above 0.95. When considering the NCBI/simNCBI database, we found that Kraken2 significantly outperformed minimap2 in mean F1 score (Figure 5C, Dunn’s test with Šidák correction for multiple comparisons), but Emu and Dnabarcoder could not be distinguished from each other, nor from either Kraken2 or minimap2. ML classifiers continued to underperform when trained on the simNCBI database, with no one *Candida* group species classified with an F1 score above 0.65 for any of the three methods tested. In a diagnostic setting, the impact of detecting a species which is not present is as important a consideration as not detecting a species which is present. To assess the behaviour of the alignment-based classifiers when a close relative of the target pathogen did not have a correct species-level assignment in the database, we determined the closest relative of our target pathogen which was present in our mock community. Based on published phylogenies (Butler et al., 2009; Tsui et al., 2008) and pairwise similarity between GSD reference sequences for *Candida* species, we determined this to be *C. dubliniensis,* which shared 92.2% sequence identity with our *C. tropicalis* GSD reference sequence. We removed the reference sequence for *C. dubliniensis* from the NCBI database, but kept query sequences from this species in the Mock 3 community, which we hypothesised would increase the false positive count for *C. tropicalis*, as *C. dubliniensis* sequences would be classified to this species. However, no significant increase in false positives for *C. tropicalis* was observed, with the maximum false positive count only two total sequences for any given classifier. Emu was the only classifier with zero false positives for *C. tropicalis* when *C. dubliniensis* was removed from the database. Classifier mean F1 scores were indistinguishable by Dunn’s test after correcting for multiple comparisons (Figure 5D), however Kraken2 and Dnabarcoder had empirically higher means, and less variability in per-species F1 score.

**Figure 5.**
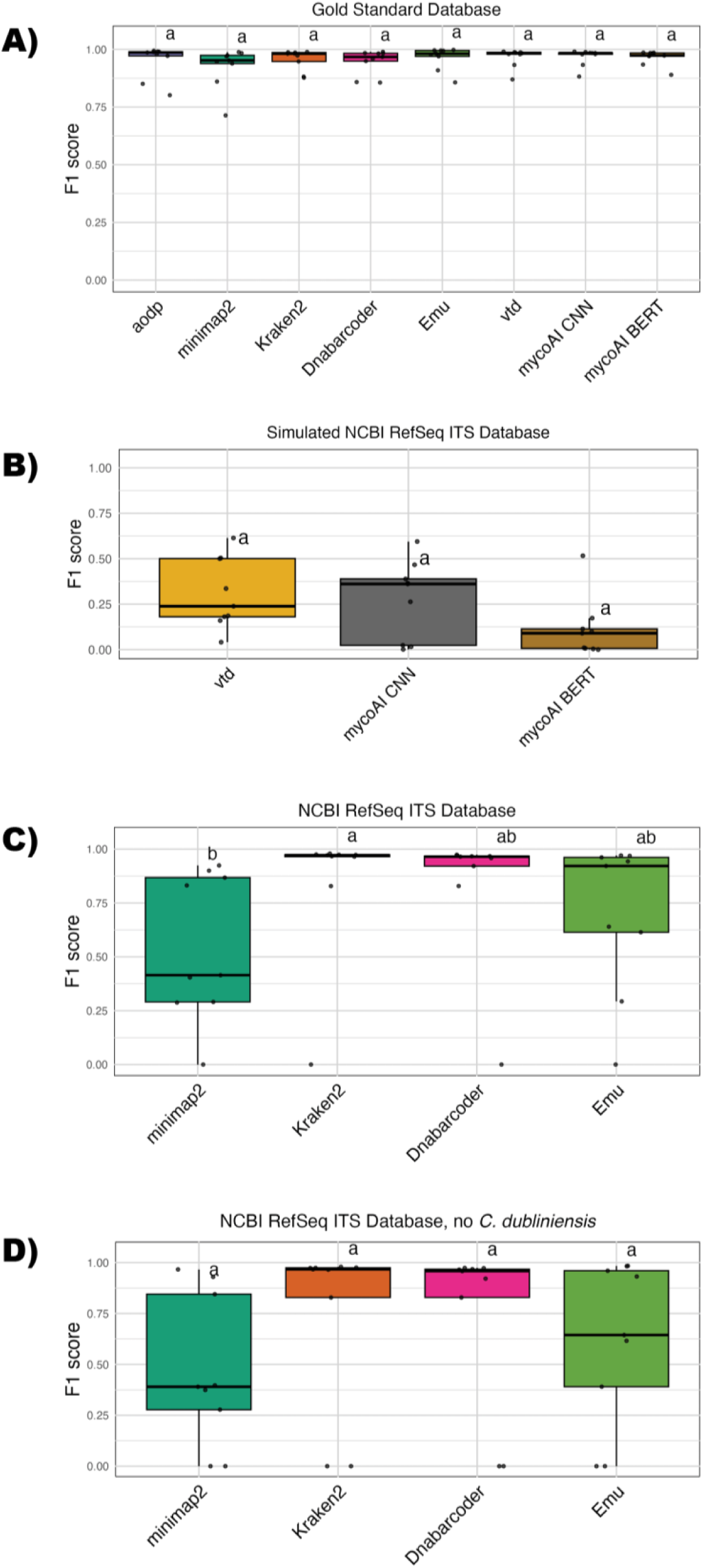
Kraken2, Dnabarcoder and Emu can control false positive and false negative classifications of target pathogen *Candida tropicalis* from amongst its close relatives. CLD letters indicate significant pairwise comparisons assessed by Dunn’s tests with Šidák correction for multiple comparisons. **A)** Classifiers can differentiate well between nine *Candida* group species using the GSD. **B)** Machine learning approaches trained on the simNCBI dataset did not perform well, consistent with previous results for this training set. **C)** Kraken2 best distinguished between *Candida* group species, but all classifiers struggled to identify *C. metapsilosis* using the NCBI RefSeq database. **D)** Removing a close relative of the target pathogen from the reference database did not result in significantly higher false positive calls for the target pathogen, but did decrease mean F1 score for minimap2 and Emu classifiers.

### 3.7 Fate of sequences from ‘unknown’ query species

We investigated the way classifiers treated sequences from ‘unknown’ origin by looking at the fate of sequences from species which did not have a correct species-level taxonomic assignment in the NCBI database, using the species in our Mock 2 community which did not appear in the NCBI RefSeq ITS database. We assessed the classification result for each species by assigning ‘closeness score’, which increased in value with every taxonomic rank between the lowest rank shared by the correct origin species, and the taxon to which the most sequences had been assigned (referred to as the top hit for that input species). For the purposes of this analysis, we removed the species for which there was a correct species-level assignment in the NCBI RefSeq ITS database.

To facilitate comparisons across alignment-based, *k-*mer and ML approaches, we compared the delta value between the lowest possible closeness score available in the appropriate database or training set, and the observed closeness score, using the NCBI Taxonomy (Schoch et al., 2020; Sayers et al., 2019) as a reference. For the evenly distributed Mock 1 community, Kraken2, Dnabarcoder and Emu were distinct from minimap2 (Figure 6A; Dunn’s tests with Šidák correction for multiple comparisons, p < 0.05). In the Mock 2 community, only Kraken2 and Dnabarcoder were distinct from minimap2 (Dunn’s test with Šidák correction, p < 0.05) (Figure 6B). These results indicated that minimap2 placed sequences from unseen species into taxonomically more distant taxa. Emu did not perform as well with the unevenly distributed community.

**Figure 6.**
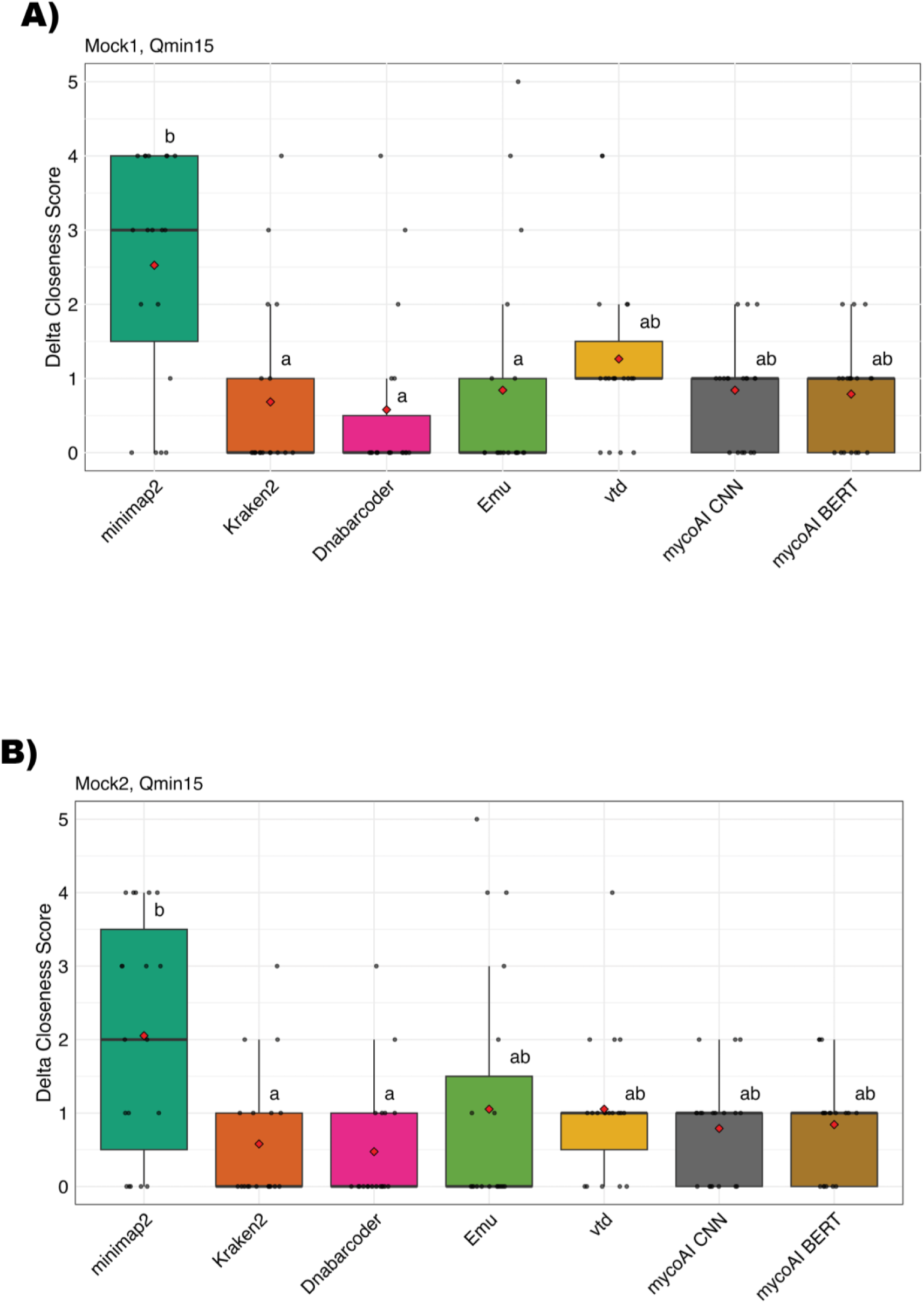
Kraken2 and Dnabarcoder best mitigate the effects of incomplete reference databases by classifying sequences from ‘unknown’ species to taxonomically closer relatives. Delta closeness score assesses the difference between the correct taxonomy for an ‘unknown’ species, and the closest possible taxonomic classification in the reference database, with each rank of difference increasing the closeness score value by one. Lower delta closeness scores indicate a classifier’s ability to place a sequence from a species with no correct species-level classification into a taxonomically closer relative. CLD letters indicate significant pairwise comparisons as determined by Dunn’s tests with Šidák correction for multiple comparisons (p < 0.05). **A)** minimap2 had a significantly higher mean delta closeness score than other alignment- or *k*-mer based approaches when assessing evenly distributed ‘unknown’ species. **B)** Kraken2 and Dnabarcoder continue to outperform other methods, with statistically distinct mean delta closeness scores in comparison to minimap2, which had a significantly higher delta closeness score in comparison to other methods.

For alignment-based and *k-*mer classifiers, we modelled the relative proportion of variance in closeness score attributable to the factors “Classifier”, “Species”, “MinSeqQual”, and number of sequences classified into the representative result taxon (“SeqsClassified”), and found that a cumulative link mixed model (CLMM) fit the data best, based on Aikake Information Criterion (AIC) value assessment. For both Mock1 and Mock2 data, minimum sequence quality did not contribute significantly to variation in classifier closeness score, with CLMMs indicating better fit (lower AIC value) when the “MinSeqQual” term was not included as a factor (ΔAIC(Mock1) = -1.86, ΔAIC(Mock2) = -1.99).

For Mock2, the more realistic, unevenly distributed community, the model which best fit the closeness score data had Classifier as a fixed factor, and Species as a random factor, indicating again that there are species-specific effects determining the performance of a given classification algorithm. This model also reinforced that minimap2 had a significant, positive effect on closeness score. The best-fitting CLMM for Mock2 had Species included as a random effect (AIC = 216.36), in comparison to an ordinal logistic regression (OLR) model which did not consider Species as a factor (AIC = 233.51). This indicates that while Species does influence closeness score variance, the contribution to variance is not consistent across distinct input Species.

### 3.8 Simulating classifier detection limits from experimental data

We determined the minimum number of sequences required for each classification method to detect each of the 54 species in our mock community from the outputs of each classifier when using the GSD. We assessed the minimum required sequencing depth at two thresholds of species abundance: detection, meaning at least one sequence assigned to all species, and minimum 109 sequences per species, meaning all species detected at ≥0.1% read abundance, relative to the total number of input sequences. We also ran the same simulation using the ground truth community abundance distribution, as a comparison to an idealised ‘perfect’ classifier.

The number of sequences required to detect all species with at least one sequence using the ‘ideal’ classifier for Mock 1 was 373 sequences. All classifiers performed similarly in this test, with all minimum sampling depths between 372 (mycoAI CNN and vtd) and 420 (aodp) (Figure 7A). When we applied the same function to Mock 2 results, where taxa are unevenly distributed, the minimum number of sequences required to detect all 54 species with at least one sequence for the idealised classifier increased to 554 (Figure 7B). This represents a ∼1.5X increase in sequences required to meet the minimum abundance threshold. We found that Dnabarcoder and the machine learning methods were most sensitive, requiring the fewest sequences to detect all species with a minimum of one sequence per species.

**Figure 7.**
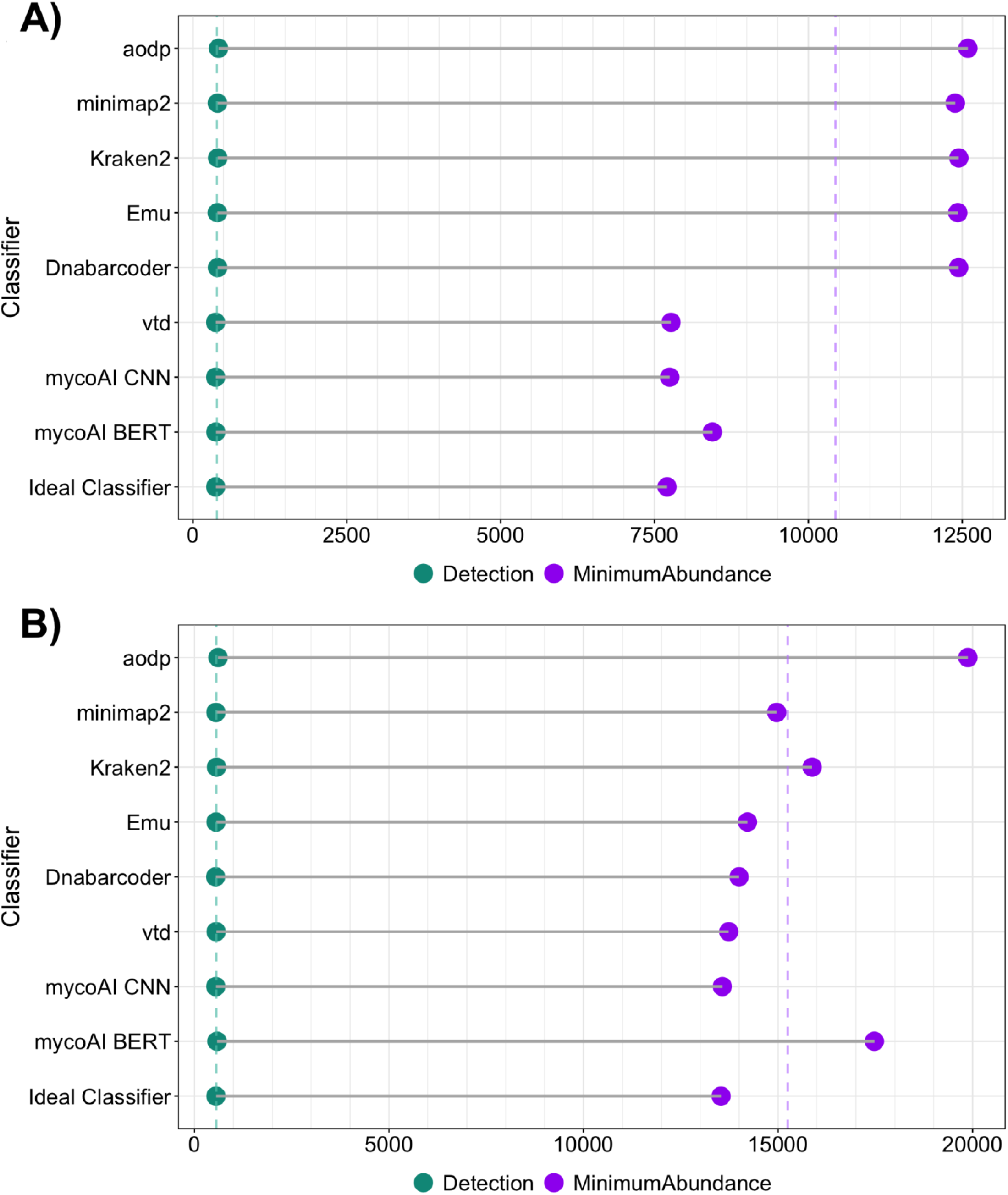
Machine learning classifiers require the lowest sequencing depth to detect all mock community species, for both even Mock 1 (A) and uneven Mock 2 (B) communities. Dotted vertical lines indicate overall mean minimum sequences required to meet abundance thresholds for all 54 mock community species across all classifiers. The ‘Ideal’ classifier results were simulated from ground truth data, and represent a classifier which perfectly reproduces community abundance distribution.

In a realistic metabarcoding study, a single sequence is often not sufficient evidence to suggest that a species is truly present. We therefore also calculated the minimum number of sequences required to detect all 54 species at a minimum of 109 sequences/species, which is equivalent to 0.1% of total input sequences. At the higher minimum abundance threshold of 109 sequences, the Mock 1 idealised classifier required 7,705 sequences to detect all species at or above this threshold. The machine learning methods performed particularly well for Mock 1, with all three models requiring the fewest sequences to detect all species (Figure 7A). In comparison, the Mock 2 idealised classifier required 13,533 sequences to detect all species at this higher abundance threshold. The mycoAI CNN model continued to perform well for this task, along with Dnabarcoder and the other CNN model, vtd (Figure 7B).

Despite Kraken2 appearing to be one of the more sensitive classifiers tested when considering the results of other tests in this benchmarking work, it was unable to resolve all taxa at the species level when classifying against the GSD. In our GSD, the reference sequences for *Botrytis cinerea* and *Botrytis fabae* are distinguished by a ∼200 bp insertion in the *B. fabae* reference sequence, but otherwise share complete sequence identity. This means that while *B. fabae* could be differentiated from *B. cinerea* using Kraken2, *B. cinerea* could not be differentiated from *B. fabae*. Because of this, there is no point at which Kraken2 would detect all species at any abundance. To overcome this, we ran an adjusted version of the function, calculating minimum abundance thresholds for 53 species.

## 4. Discussion

We benchmarked eight classification approaches spanning alignment algorithms (minimap2, Emu, Dnabarcoder), *k-*mer detection methods (Kraken2, aodp), and machine learning algorithms (mycoAI CNN and BERT, ‘vtd’ CNN) using an *in silico* mock community of real ONT long-read fungal metabarcodes. The sequences in our mock community were generated using a ‘universal’ ITS primer pair which amplifies not only both ITS1 and ITS2, but also several hundred base pairs of both the 18S SSU and 28S LSU genes (White et al., 1990). Increased metabarcode sequence length provides more comparative information when making taxonomic assignments, and has improved metabarcode classification to the species, or even strain level, in bacteria (Johnson et al., 2019; Matsuo et al., 2021), and invertebrates (Baloğlu et al., 2021; Krehenwinkel et al., 2019). While ONT sequencing has shown promising results for metabarcoding of Fungi (Lu et al., 2022; Hu et al., 2022; D’Adreano et al., 2021), this work is the first comprehensive benchmark specifically applying ONT long-read metabarcoding to a fungal mock community, assessing classification at the species level.

While our benchmarking work addresses the common influential factors in classification of fungal metabarcodes such as classifier and database choice, we wanted to address this from the context of two broad use cases for metabarcoding – biodiversity assessment, and diagnostics. These use cases are reflected in our database choices. The gold standard database (GSD) represents a small, targeted database of high-quality reference sequences, where each species in the mock community is expected to be represented. These smaller databases have demonstrated improved classification of metabarcodes in diagnostic applications (Irinyi et al., 2015; Langsiri et al., 2023; Ohta et al., 2023; Chen et al., 2022), and in biodiversity assessment of underexplored ecosystems (Philip et al., 2024). In comparison, we chose the NCBI RefSeq Targeted Loci ITS database (O’Leary et al., 2016; Schoch et al., 2014) as an example of a publicly available reference database which may be used for biodiversity assessment in a microbiome study. While the NCBI ITS database does not contain reference sequences for as many fungal species as others such as UNITE (Abarenkov et al., 2024) or GlobalFungi (Větrovský et al., 2020), it contains significantly more than our GSD.

### 4.1 Classifier choice can mitigate the influence of incomplete reference databases

We assessed classifier performance based on three measures of community composition: F1 score, alpha diversity metric Shannon index, and Bhattacharyya distance. In our analyses, we found that false negative classifications were more frequent and variable than false positive classifications – that is, that mock community species were more likely to be missed, than be falsely detected. This likely contributed to our observation that “Species” was more important than “Classifier” in determining F1 score in our random forest modelling, both when using the GSD, and when using the NCBI database as a reference. One of the commonalities between these databases is that species were represented by very few reference sequences. Despite our GSD reference sequences being built from the same ITS amplicons as the mock community query sequences, representing each species with a single consensus sequence limits the amount of intraspecific variation which can be expressed in the database. This can lead to oversplitting, or unclassified sequences, when sequence variation exceeds similarity thresholds for determining matches between query and reference sequences (Nilsson et al., 2008; Bradshaw et al., 2023).

While we note that these data are the result of classification against a single reference database, it is a well-established critique of metabarcoding, particularly in its application to microbiome analyses, that reference databases are a primary limitation to accurate data interpretation (Furneaux et al., 2021; Keck et al., 2023; Abarenkov et al., 2022; Kidd et al., 2023; O’Donnell et al., 2015). Addressing this limitation would require enormous investment of time and resources, given the diversity of fungal taxa, but this does not mean that the effect of incomplete databases cannot be mitigated and minimised with informed experimental design. Our results indicate that classifiers such as Dnabarcoder, Kraken2, and under some conditions, machine learning methods such as the mycoAI CNN model, are suited to address this limitation, as they create informed discriminatory thresholds at the species level based on the reference database itself. This is why these classifiers could better place ‘unknown’ taxa into taxonomically close relatives: species-level discrimination is based on what is actually possible given the interspecific variation represented in the reference database, rather than applying predetermined thresholds which may not accurately represent species boundaries. This is an important consideration in biodiversity assessment applications using eDNA, where species composition is not necessarily known.

While species-level identification may not be possible while reference databases are incomplete, classifier choice can improve the confidence of classification results at higher taxonomic levels such as genus and family. Based on our results in both eDNA diversity and diagnostic applications, Dnabarcoder appears to be the most suitable classification approach for long-read ITS metabarcoding of Fungi. Kraken2 and CNN models may also be candidates in some cases, provided that users are aware of their caveats regarding database or training set requirements.

### 4.2 Balancing sequence quality and sequence length in metabarcoding

A consistent observation across our statistical analyses and modelling, was that the factor ‘Minimum Sequence Quality’ did not significantly influence the variation in any classification performance metrics tested. In this benchmarking study we filtered sequences to a minimum sequence Phred score of 15, which is approximately 96.8% mean sequence accuracy, and 17, which is approximately 98% mean accuracy. In comparison, Illumina sequencing instruments have a mean sequence accuracy of >Q30, (mean accuracy above 99.9%), which we appreciate is still higher than the quality filtering thresholds used in this study. There are applications where the accuracy of Illumina sequences makes it a more suitable technology, but we argue that equally, there are applications where the resolution possible with ONT long-read sequences may be a higher priority. Whether in a diagnostic case, where species-level resolution could inform therapeutic decision-making, or biodiversity assessment, where accurate species richness estimations may inform policy or funding, accurate species detection in these situations draws conclusions which could not be reached with genus-level data alone. Therefore, while we acknowledge that ONT long-read data does not yet reach the same sequence accuracy as Illumina instruments with consistency, the comparative information gained with greater sequence length should factor into decision making during experimental design of fungal metabarcoding studies. This design will then need to inform classification approach, as different algorithms, query sequence lengths and reference databases will influence the way classifiers interpret interspecific variation, and therefore how sequences are classified. Using informed similarity cut-offs based on the users’ experimental design will best improve ITS metabarcode classification, particularly from eDNA samples.

### 4.3 Future perspectives for machine learning classifiers in fungal metabarcoding

Machine learning methods are one of the most recent approaches to DNA sequence classification. In our benchmark, we tested two CNN models, ‘vtd’ and the mycoAI CNN model, and one BERT model, also from mycoAI. These models performed exceptionally well when trained on sequences from the GSD, returning the highest F1 scores, lowest Bhattacharyya distances, and requiring the fewest sequences to detect all species at both simulated minimum abundance thresholds. Together, this suggests that not only are these models accurately reproducing fungal community compositions, but that they are also highly sensitive, making them suited to diagnostic tasks as well. Overall, these models were also robust to classify species which were not represented in the training data. ML methods demonstrated low delta closeness scores, which indicated that they were able to place ‘unknown’ species taxonomically closer to their true origin. The vtd CNN classifier struggled to correctly place *Fusarium proliferatum*, and *Scedosporium boydii*, for both of which there was a genus-level match in the training data. This was one of the few times that minimum sequence quality did influence classifier performance, as F1 scores for these species improved for this classifier when tested with Qmin17 data, as opposed to Qmin15 data (Supplementary Table T2). However, both mycoAI models were able to correctly place sequences from these species to their genus-level matches at the lower Qmin15 sequence quality threshold.

The caveat, and clear next step for increasing adoption of ML methods for sequence classification, is the requirement for more complex training sets than current reference databases provide. A part of this challenge is that ML models do not represent sequence similarity in a way that it is easily human interpretable. Pairwise sequence similarity, the way it is assessed by *k*-mer and alignment classifiers, is a very intuitive way to consider taxonomic classification. It is directly linked to sequence evolution – the more diverged two species are, the less sequence similarity they would be expected to share. ML methods do assess sequence similarity, but in a more abstract way. ML models take a (labelled) training set, and learn the features which distinguish different classes directly from the training data. For DNA sequence classification, this can include short subsequences, similarly to *k*-mer methods, which are distinctive for different species, but can also include hallmarks such as sequence length. Similarly to reference databases used for canonical alignment or *k*-mer based approaches, how well the training set represents true sequence diversity will significantly contribute to the fitness of a machine-learning model.

The number of sequences per species for adequate model training depends on the amount of variation in the test set. This is what makes most available reference databases unsuitable for direct use as a training set for ML methods. For many DNA barcode reference databases, the distribution of sequences per species is skewed, with overrepresentation of well-studied organisms, and many underrepresented, or uncharacterised species (Kvist et al., 2013; Fort et al., 2022; Keck et al., 2023). For models like neural networks, such an unbalanced training set could compromise classifier performance. The length distribution of the training set will also influence how well-trained the model is. For supervised learning methods, features such as sequence length will be learned by the model and used to discriminate between species. If the training set is not representative of the length distribution of the query sequences (test set), then these features will not be informative. Additionally, if the training set sequences for a single species have a broad length distribution, this will also make sequence length an uninformative feature, as there is no distinct feature for the model to learn for this species.

The models we trained on the GSD training set were well-trained without overfitting using 2,200 sequences/species (Supplementary File S7), which is only achievable for a fraction of characterised fungal species in reference databases. Until more comprehensive training sets are available at the species level, ML classifiers could be used to classify sequence data to higher taxonomic levels, such as family, and other classification algorithms used to distinguish at the genus and/or species level.

### 4.4 Metabarcoding for fungal diagnostics

One of the use cases we used as a context for our benchmarking experiments was diagnosis of fungal infections with ITS metabarcodes. For this use case, we wanted to emphasise diagnostic priorities – limiting false positives as well as false negatives, minimum detection thresholds, and confident discrimination between closely related species. To do this, we created ‘Mock 3’, which contained all taxa present in the previous mock communities, however the analysis was limited to the correct identification and abundance estimation of the clinically relevant *Candida* group, focusing on pathogenic species *Candida tropicalis*. *C. tropicalis* is a causative agent of superficial and systemic Candidaemia in humans, and is increasingly reported as resistant to many first-line antifungals (dos Santos and Ishida, 2023; Zuza-Alvez et. al., 2017). Additionally, it is closely related to other *Candida* group members in the mock community (Tsui et. al., 2008), sharing high nucleotide identity at the ITS loci. Given that classification methods did not struggle to delineate between most *Candida* group members when using the GSD, or the full NCBI database, we introduced another degree of complexity by removing a close relative. When we removed *C. dubliniensis* from the NCBI database, we hypothesised that *C. tropicalis* abundance would be falsely inflated due to false positive classifications of sequences originating from *C. dubliniensis.* However, we found that alignment-based classifiers did not significantly overestimate *C. tropicalis* abundance. While there are closer relatives to *C. dubliniensis* than *C. tropicalis*, such as *C. albicans*, it was surprising to see no significant increase in false positive calls for *C. tropicalis*. Extending this experiment to include other species groups, particularly from pathogens which are traditionally difficult to delineate at the species level, such as members of the *Aspergillus* species complex, would be a worthwhile investigation for applying this method for diagnostic purposes.

A species which classifiers did struggle to distinguish in the Mock 3 experiment was *C. metapsilosis*. *C. metapsilosis* is considered a part of the *Candida parapsilosis* complex, containing *C. parapsilosis* sensu stricto, *C. metapsilosis*, and *C. orthopsilosis* (Govrins and Lass-Flörl, 2024), all of which are pathogenic to humans (Bonfietti et al., 2012; Feng et al., 2011). The reference ITS sequences for *C. metapsilosis* and *C. orthopsilosis* contained in the NCBI RefSeq database share 94.8% nucleotide identity, and no alignment or *k*-mer based classifier was able to correctly classify *C. metapsilosis* using this database. Interestingly, the two CNN classifiers which were trained on the simNCBI database correctly determined a small number of sequences as *C. metapsilosis* (Supplementary Table T3). The simNCBI training set contained long ITS sequences reconstructed from reference sequences available in the NCBI RefSeq ITS, 28S and 18S databases, though of these, *C. metapsilosis* reference sequences were only available in ITS and 28S RefSeq databases. While overall, this training set performed poorly, the ability of machine learning methods to correctly classify any *C. metapsilosis* query sequences is an improvement from alignment and *k*-mer based methods, which could not classify any sequences as *C. metapsilosis* when using the NCBI RefSeq ITS database. This demonstrates the power of machine learning approaches for highly sensitive classification of fungal ITS sequences. It also demonstrates the applicability of long-read ITS sequencing in diagnostic use cases, as some sequence features which allowed machine learning methods to classify *C. metapsilosis* sequences correctly may be contained in the 28S locus, a region which was not represented in the NCBI database used with alignment and *k*-mer approaches.

It is imperative to understand the detection limit of any diagnostic method, as this is a measure of the sensitivity of the approach. This is exacerbated in ONT metabarcoding applications, because unlike Illumina’s sequencing-by-synthesis technology, ONT sequencers are limited by the number of DNA molecules in the prepared library. In samples with low-abundance species, either due to extraction biases or low growth rate, understanding how classifier choice may influence detection thresholds informs minimum required sequencing depth, and therefore cost-effectiveness of the method. We used our real classification results to calculate the minimum number of sequences required to classify all 54 species in our mock communities based on a probabilistic Monte Carlo simulation. By calculating the probability of detecting all 54 species together, we can model the cumulative effect of the variation introduced by the classification method on the total mock community. The underlying species distribution then further influences this calculation, as the more uneven the community abundance distribution is, the more sequences will be required to detect rare species at minimum abundance thresholds.

## 5. Conclusions

In this work, we present a benchmark of classification methodologies for ONT long-read metabarcodes of fungi. Our results demonstrate classifiers such as Kraken2 and Dnabarcoder which determine informed thresholds of sequence similarity outperform other classification approaches in terms of precision, faithfully reproducing species abundance distributions, sensitivity, and ability to appropriately handle unseen species. This is particularly the case when long-read ITS metabarcodes are classified against a reference database of shorter ITS sequences, such as those which populate most publicly available ITS databases. Our work also indicates that machine-learning approaches hold great promise when provided a representative training dataset, however, this caveat is largely not met by currently available reference databases. To better leverage the potential of ML classifiers, more comprehensive, and representative, training datasets are required.

## Data availability

All input sequence data used in this benchmarking work can be accessed from Zenodo: 10.5281/zenodo.15751793

All scripts used to quality control, analyse, and generate figures for this benchmarking work can be accessed from the associated GitHub repository: https://github.com/AbiGraetz/Nanopore-ITS-Benchmarking

