## Supplementary Files for "Benchmarking fungal species classification using Oxford Nanopore Technologies long-read ITS metabarcodes"

**Supplementary File S1 – *In silico* mock community composition and species abundance**

**Supplementary File S2 – PCR and sequencing library preparation protocol for long-read fungal ITS metabarcoding**

**Supplementary File S3 – Sequence quality control workflow**

**Supplementary File S4 – Gold Standard Database consensus sequence building**

**Supplementary File S5 – Classification F1 score for machine learning classifiers using the NCBI RefSeq ITS database**

**Supplementary File S6 – Precision and recall rate for classification approaches using the GSD**

**Supplementary File S7 – Machine learning model overfitting**

| **Supplementary File S1 -** Mock community species abundances in each mock community, and mean sequence length after quality control. Mean sequence length calculated based on the sequences selected for Mock1. | | | | |
| --- | --- | --- | --- | --- |
| **Species** | **Abundance (sequences) Mock1** | **Abundance (sequences) Mock2** | **Abundance (sequences) Mock3** | **Mean sequence length (+/- SD)** |
| [Candida] *boleticola* | 2000 | 1090 | 1514 | 2109.5 bp  (+/- 487.29) |
| [Candida] *caryicola* | 2000 | 1090 | 1215 | 1978.0 bp  (+/- 383.12) |
| *Diutina catenulata* | 2000 | 1135 | 1364 | 1858.7 bp  (+/- 311.74) |
| *Candida dubliniensis* | 2000 | 1090 | 1816 | 2102.7 bp  (+/- 381.68) |
| *Nakaseomyces glabratus* | 2000 | 2729 | 1971 | 2259.5 bp  (+/- 554.17) |
| *Candida metapsilosis* | 2000 | 1983 | 121 | 2096.6 bp  (+/- 333.34) |
| *Candida parapsilosis* | 2000 | 3182 | 1891 | 2091.6 bp  (+/- 359.64) |
| *Candida tropicalis* | 2000 | 2281 | 840 | 2066.3 bp  (+/- 389.17) |
| [Candida] *zeylanoides* | 2000 | 1494 | 926 | 2216.8 bp  (+/- 326.07) |
| *Puccinia striiformis* f. sp. *tritici* | 2000 | 1205 | 1494 | 2279.8 bp  (+/- 369.34) |
| *Puccinia triticina* | 2000 | 1565 | 2785 | 2279.6 bp  (+/- 422.42) |
| *Puccinia graminis* f. sp. *tritici* | 2000 | 3514 | 1639 | 2300.2 bp  (+/- 362.08) |
| *Leptosphaeria maculans* | 2000 | 1957 | 2934 | 2154.8 bp  (+/- 311.83) |
| *Sclerotinia sclerotiorum* | 2000 | 3685 | 2564 | 2420.0 bp  (+/- 387.52) |
| *Botrytis cinerea* | 2000 | 2318 | 2058 | 2142.3 bp  (+/- 289.09) |
| *Botrytis fabae* | 2000 | 1090 | 1901 | 2399.4 bp  (+/- 411.03) |
| *Zymoseptoria tritici* | 2000 | 1090 | 1545 | 2124.3 bp  (+/- 335.73) |
| *Sporisorium scitamineum* | 2000 | 1330 | 2819 | 2272.0 bp  (+/- 441.01) |
| *Erysiphe necator* | 2000 | 3150 | 2383 | 2049.6 bp  (+/- 580.86) |
| *Austropuccinia psidii* | 2000 | 1893 | 2229 | 2282.6 bp  (+/- 375.69) |
| *Pyrenophora tritici-repentis* | 2000 | 2253 | 2731 | 2196.0 bp  (+/- 410.45) |
| *Cryptovalsa ampelina* | 2000 | 1937 | 2006 | 2169.3 bp  (+/- 367.30) |
| *Eutypa lata* | 2000 | 1210 | 2728 | 2847.5 bp  (+/- 552.51) |
| *Diplodia seriata* | 2000 | 3763 | 2866 | 2160.9 bp  (+/- 349.96) |
| *Fusarium pseudograminearum* | 2000 | 2358 | 2218 | 2138.9 bp  (+/- 296.21) |
| *Aspergillus flavus* | 2000 | 1090 | 2069 | 1804.2 bp  (+/- 726.35) |
| *Aspergillus fumigatus* | 2000 | 2580 | 1489 | 1610.1 bp  (+/- 728.27) |
| *Blastobotrys adeninivorans* | 2000 | 1301 | 2949 | 2136.2 bp  (+/- 373.58) |
| *Blastobotrys proliferans* | 2000 | 1090 | 1361 | 2124.1 bp  (+/- 312.3) |
| *Cryptococcus gattii* VG I | 2000 | 2791 | 1740 | 2035.0 bp  (+/- 548.81) |
| *Cryptococcus neoformans* VN IIII | 2000 | 2711 | 1746 | 2142.7 bp  (+/- 383.23) |
| *Fusarium proliferatum* | 2000 | 2566 | 2238 | 2098.8 bp  (+/- 386.59) |
| *Geotrichum candidum* | 2000 | 2419 | 2519 | 1818.4 bp  (+/- 313.9) |
| *Kluyveromyces lactis* | 2000 | 1748 | 2787 | 2095.9 bp  (+/- 383.7) |
| *Kluyveromyces marxianus* | 2000 | 1483 | 2508 | 2228.6 bp  (+/- 424.03) |
| *Kodamaea ohmeri* | 2000 | 1401 | 2785 | 1882.4 bp  (+/- 352.01) |
| *Penicillium chrysogenum* | 2000 | 1589 | 2276 | 2113.6 bp  (+/- 457.20) |
| *Pichia kudriavzevii* | 2000 | 1090 | 2709 | 2087.6 bp  (+/- 348.95) |
| *Pichia membranifaciens* | 2000 | 4179 | 2720 | 1695.5 bp  (+/- 669.04) |
| *Rhodotorula mucilaginosa* | 2000 | 3132 | 1933 | 2189.3 bp  (+/- 361.10) |
| *Scedosporium aurantiacum* | 2000 | 1090 | 1282 | 2811.7 bp  (+/- 713.07) |
| *Scedosporium boydii* | 2000 | 1377 | 2887 | 2153.5 bp  (+/- 416.36) |
| *Trichomonascus ciferrii* | 2000 | 1307 | 1942 | 2126.1 bp  (+/- 445.95) |
| *Trichosporon asahii* | 2000 | 2666 | 2027 | 2118.4 bp  (+/- 406.08) |
| *Wickerhamomyces anomalus* | 2000 | 1725 | 1282 | 2148.8 bp  (+/- 429.85) |
| *Yamadazyma mexicana* | 2000 | 2092 | 1517 | 2177.9 bp  (+/- 324.22) |
| *Yamadazyma scolyti* | 2000 | 1785 | 1533 | 2193.2 bp  (+/- 293.22) |
| *Yarrowia lipolytica* | 2000 | 1769 | 1328 | 1884.6 bp  (+/- 256.39) |
| *Diaporthe* sp. CCL067 | 2000 | 3307 | 2495 | 2160.6 bp  (+/- 353.89) |
| *Quambalaria cyanescens* CCL055 | 2000 | 1570 | 2368 | 2074.3 bp  (+/- 620.53) |
| *Entoleuca* sp. CCL052 | 2000 | 3468 | 1819 | 2506.9 bp  (+/- 467.87) |
| *Cortinarius globuliformis* CM4 | 2000 | 1090 | 1932 | 1698.2 bp  (+/- 797.02) |
| *Asteroma* sp. CCL060 | 2000 | 1951 | 1652 | 2135.0 bp  (+/- 358.9 bp) |
| *Tuber brumale* | 2000 | 2051 | 2581 | 1952.8 bp  (+/- 903.45) |

**Supplementary File S2**

**PCR and library preparation protocols for long-read fungal ITS metabarcoding**

**Stage 1: PCR Amplification**

A PCR master mix with 50 µL reaction volume was prepared for all samples, plus an additional 10% dead volume as error allowance, in PCR 8-strip tubes, using the following reagents:

| **Component** | **Supplier** | **Stock Concentration** | **Final Concentration** | **Volume per reaction (µL)** |
| --- | --- | --- | --- | --- |
| Sterile nuclease-free water | Invitrogen | NA | NA | 31.5 |
| 5X Q5 Buffer | New England Biolabs (NEB) | 5X | 1X | 10 |
| dNTPs | NEB | 10 mM | 0.2 mM | 1 |
| Forward Primer (NS5) | Integrated DNA Technologies (IDT) | 10 µM | 0.5 µM | 2.5 |
| Reverse Primer (LR6) | IDT | 10 µM | 0.5 µM | 2.5 |
| Q5 DNA Polymerase | NEB | 2,000 U/mL | 20 U/mL | 0.5 |
|  |  |  |  | Aliquot 48 µL |

2 µL of sample DNA was added per well, and samples flick-mixed and spun down. Samples were transferred to an Eppendorf Mastercycler Nexus 2 and cycled at the following conditions:

STEP 1	98 C	30 sec	*Initial denaturation*

STEP 2	98 C	15 sec	*Denaturation*

STEP 3	78 C	10 sec	*PNA annealing*

STEP 4	58 C	30 sec	*Primer annealing*

STEP 5	72 C	2 min	*Extension*

GO TO STEP 2 for 34 CYCLES

STEP 6	72 C	2 min	*Final extension*

STEP 7	4 C	forever	*Hold*

**Stage 2: Quality-control and cleaning**

After PCR, amplification in all samples was assessed by gel electrophoresis in 1% agarose and 0.5X TBE running buffer with SYBRSafe gel dye. Samples prepared with 6X Purple loading dye (NEB), and 1 kB Plus DNA Ladder (NEB) used for reference. Gel run at 100 V for 1 hour before imaging on a GelDoc (BioRad). Concentration of samples was measured with the Qubit dsDNA broad-range assay (Invitrogen) using 1 µL of PCR product per sample.

PCR products were cleaned with fresh 70% ethanol using 1.2X AMPure XP SPRI beads (Beckman-Coulter) to remove remaining PCR reagents before sequencing library preparation. Samples eluted in 21 µL of sterile nuclease-free water.

**Stage 3: Library preparation and sequencing**

Samples were prepared for sequencing following the Oxford Nanopore Technologies protocol Ligation Sequencing Amplicons V14 (SQK-LSK114, version current as of March, 2023), using SQK-NBD114.96 native barcoding kit to uniquely barcode each sample. Library was loaded onto a MinION R10.4 flowcell, and run set up using the MinKNOW graphical user interface.

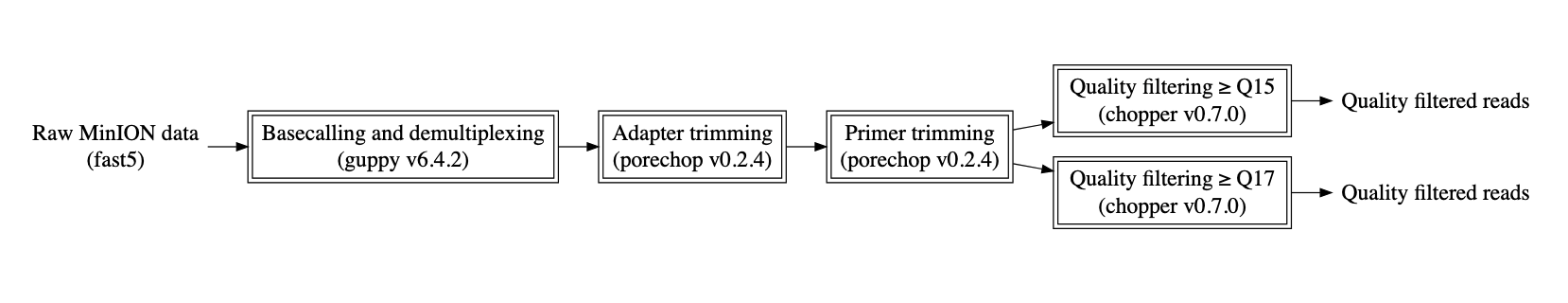

**Supplementary File S3 – Sequence quality control workflow for ONT MinION-sequenced fungal ITS am plicons.** Exact commands are available from the GitHub repository for this manuscript, as detailed in the Data Availability section.

**Supplementary File S4 – Gold Standard Database (GSD) consensus sequence building**

Consensus sequences were built from quality controlled FASTQ sequences. For each species sequenced, the sequence length between the forward and reverse primers would be a certain value, which would be variable between species. After pre-processing, we expect the length of all sequences to be approximately the length between primer binding sites.

For each species, we calculated how many sequences were in each 100 bp interval of sequence length (e.g., 100-200 bp, 200-300 bp, etc.). We used the sequences from the length ranges which the largest number of sequences, to a maximum of three discrete length ranges, to exclude outliers. We then used VSEARCH (Rognes et al., 2016) to cluster the selected sequences for each species, setting the similarity to global similarity and identity threshold to 0.95, meaning that sequences with 95% nucleotide identity would be clustered into a single group.

As we would expect the largest cluster to represent the most common ITS sequence variant for that species, the consensus sequence generated by VSEARCH for the largest cluster was used as the consensus sequence of that species. Consensus sequences generated for all 54 species were compiled into a FASTA file to use as the Gold Standard Database.

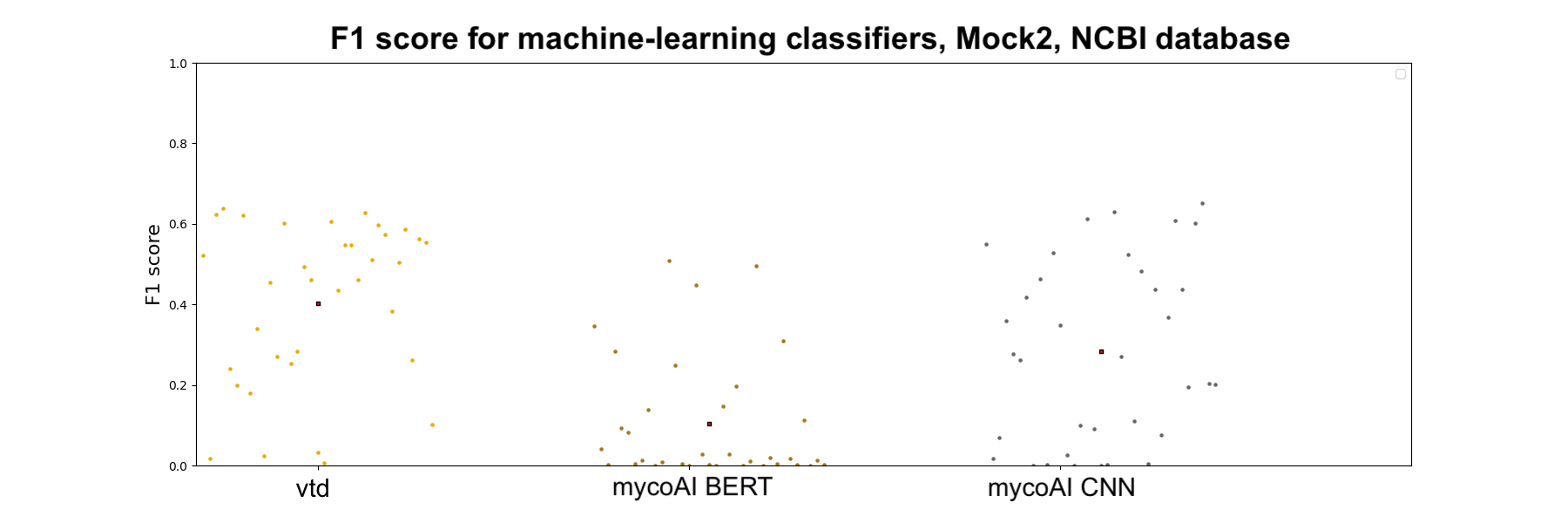

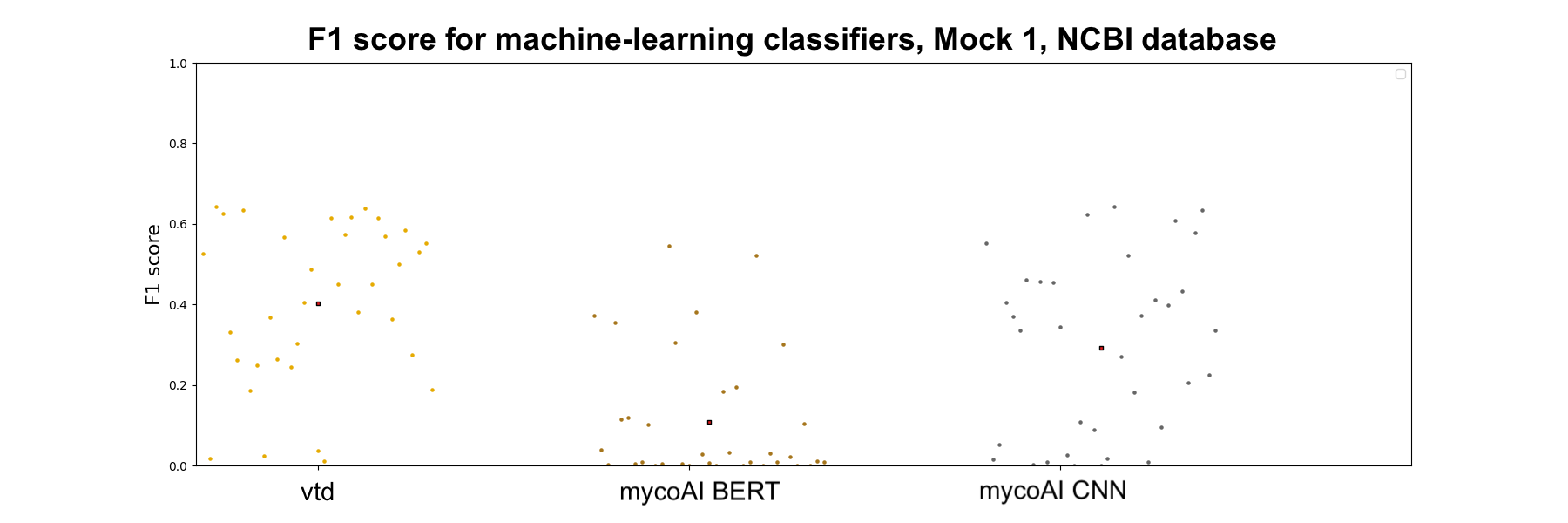

**Supplementary File S5 – F1 score for machine learning classifiers using the NCBI RefSeq ITS database.** For both Mock 1 (above) and Mock 2 (below), no classifier achieved a mean F1 score above 0.4.

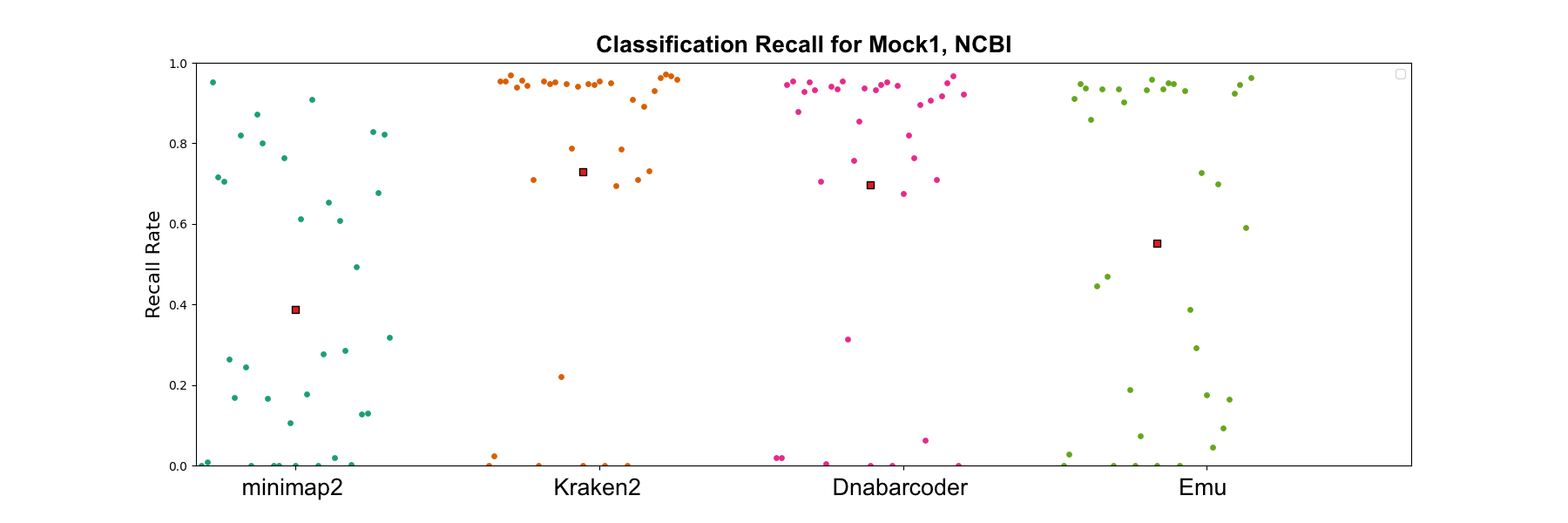

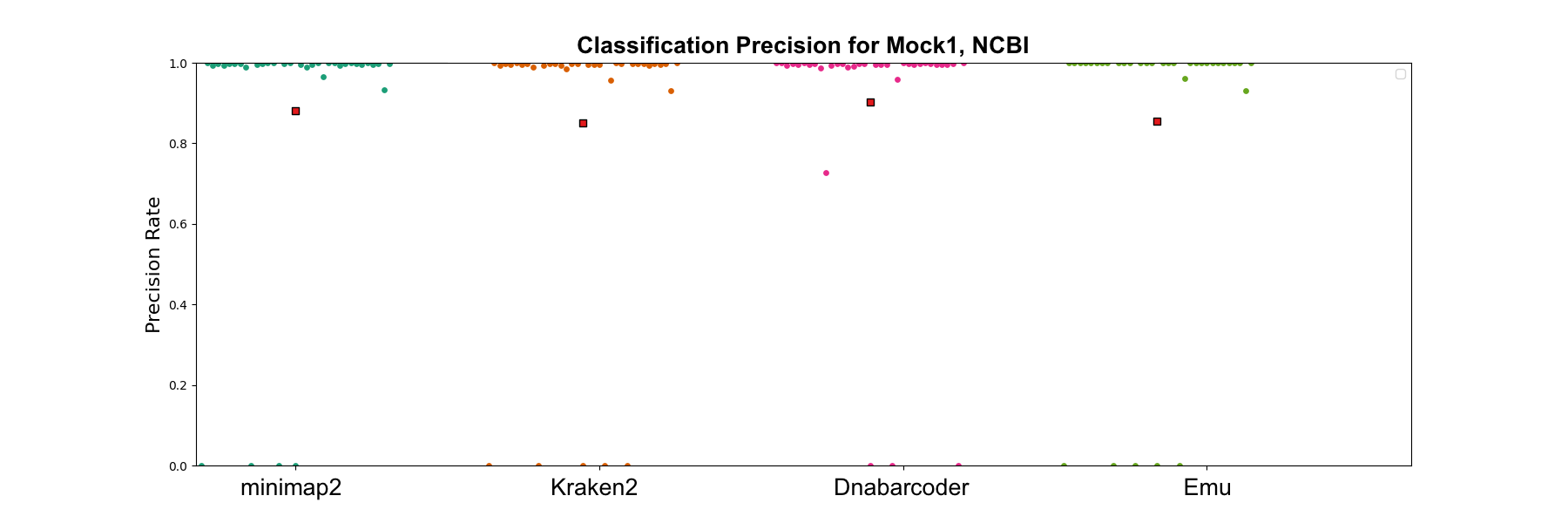

**Supplementary File S6 – Classification recall was more variable than classification precision for even Mock 1 communities when classified against the NCBI RefSeq ITS database.** This indicates that F1 score, which is calculated from the harmonic mean of precision and recall, was more influenced by variable false negative rates (captured in recall) rather than false positive rates (captured in precision).

**Supplementary File S7** - **Machine learning model overfitting**

We evaluated model overfitting using the three ML classifiers, the ‘vtd’ CNN, mycoAI CNN and mycoAI Bert, with three species as a test set. The three species represent a species with large intraspecific variation, and two with small intraspecific variation, as assessed using MAFFT (Katoh et al., 2013) and distmat (Rice et al., 2000). Species sequence variations are summarised in Supplementary Table S7.1.

| **Supplementary Table S7.1 - Mean and variance of the bases difference in the sequences of the species in the train set** | | |
| --- | --- | --- |
| Species | Mean in training set | Variation in training set |
| *Candida metapsilosis* | 13.1914 | 3230.5844 |
| *Candida parapsilosis* | 13.3896 | 3231.4510 |
| *Pichia membranifaciens* | 63.1168 | 9234.2180 |

Classification results for the three ML classifiers were saved every five epochs. Supplementary Figure S7.1 shows the number of correct sequences for the three species tested on the middle output models. From all six subfigures, there is no number of correctly classified sequences that significantly decrease after a certain number of epochs. We can also see that the models could be trained in a very small number of epochs, however, we still use the default parameters of the models. This also proves that the model itself could overcome the overfitting issue when training large number of epochs. Although there are slight shakings on each figure, that is not the result of large continuous decrease in classification accuracy after increasing to the highest point. The default batch-sizes of all the models during the training process are small, and the default learning rates could easily cause the shakings.

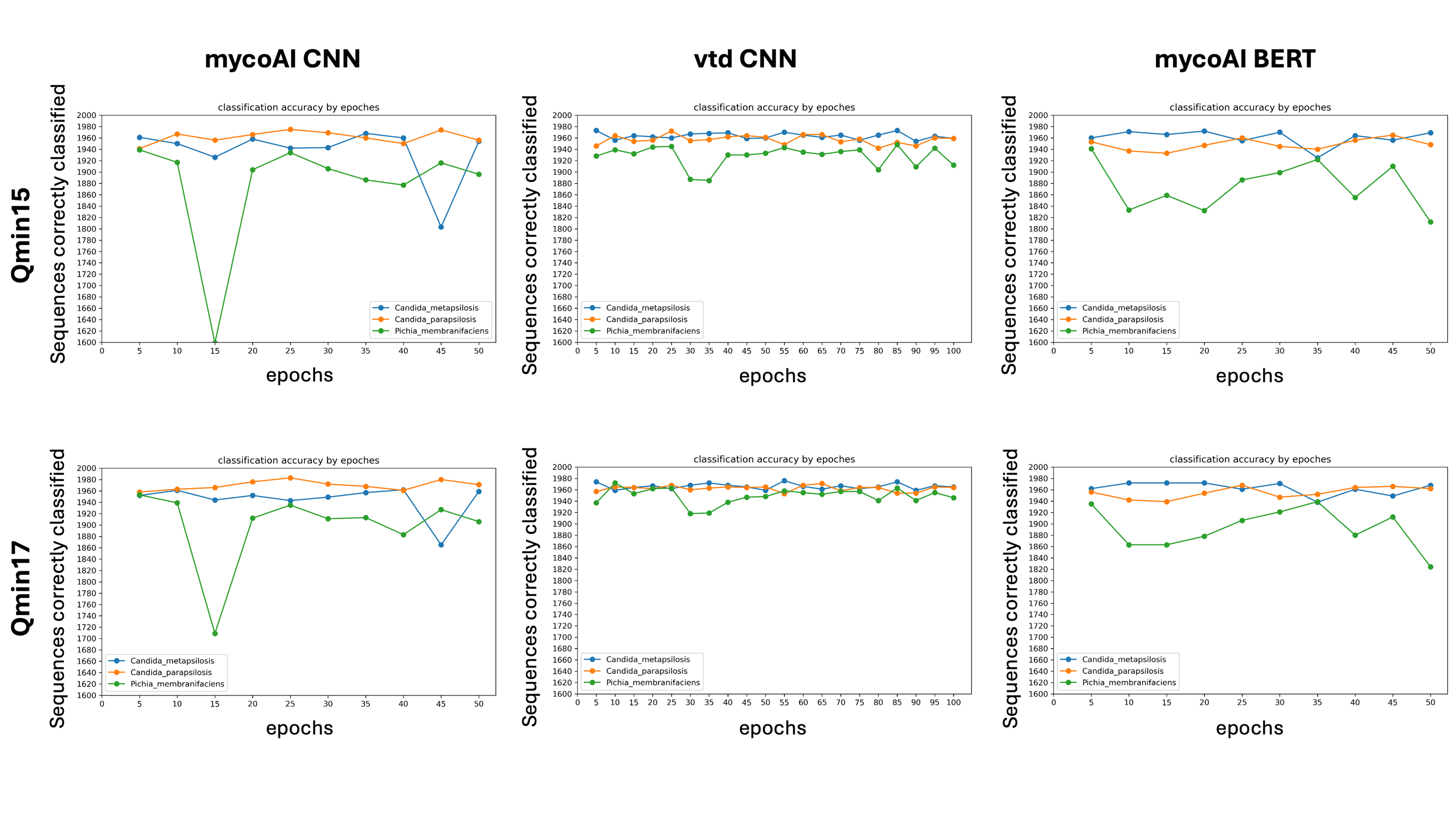

**Supplementary Figure S7.1 – Machine learning models did not demonstrate significant signs of overfitting.**
